# Diffusion kurtosis MRI tracks gray matter myelin content in the primate cerebral cortex

**DOI:** 10.1101/2024.03.08.584058

**Authors:** Colin Reveley, Frank Q Ye, David A Leopold

## Abstract

Diffusion magnetic resonance imaging (dMRI) has been widely employed to model the trajectory of myelinated fiber bundles in white matter. Increasingly, dMRI is also used to assess local tissue properties throughout the brain. In the cerebral cortex, myelin content is a critical indicator of the maturation, regional variation, and disease related degeneration of gray matter tissue. Gray matter myelination can be measured and mapped using several non-diffusion MRI strategies; however, first order diffusion statistics such as fractional anisotropy (FA) show only weak spatial correlation with cortical myelin content. Here we show that a simple higher order diffusion parameter, the mean diffusion kurtosis (MK), is strongly correlated with the laminar and regional variation of myelin in the primate cerebral cortex. We carried out ultra-high resolution, multi-shelled dMRI in ex vivo marmoset monkey brains and compared dMRI parameters from a number of higher order models (diffusion kurtosis, NODDI and MAP MRI) to the distribution of myelin obtained using histological staining, and via Magnetization Transfer Ratio MRI (MTR), a non-diffusion MRI method. In contrast to FA, MK closely matched the myelin content assessed by histology and by MTR in the same sample. The parameter maps from MAP-MRI and NODDI also showed good correspondence with cortical myelin content. The results demonstrate that dMRI can be used to assess the variation of local myelin content in the primate cortical cortex, which may be of great value for assessing tissue integrity and tracking disease in living human patients.

## Introduction

Diffusion magnetic resonance imaging (**dMRI**) of living and fixed brain tissue is increasingly used to study the cerebral cortex gray matter (Howard et al., n.d.; Wang et al., 2021), including its development (Ouyang et al., 2019; Paydar et al., 2014), areal divisions (Avram et al., 2022; Howard et al., n.d.; Liu et al., 2018), and pathology (Andica et al., 2020; McKavanagh et al., 2019; Torso et al., 2021). The way in which the cellular anatomy of the cortical gray matter shapes the dMRI signal is therefore a topic of great interest for both basic and clinical neuroimaging (Liu et al., 2018; McKavanagh et al., 2019; Torso et al., 2023; Weston et al., 2020). Human cortical gray matter is a thin (2-4 mm) corrugated sheet that envelops the vast (2000 cm^2^) cerebral surface. This surface is composed of several dozen cortical regions which vary in their cellular(Brodmann, 2006) and myelin (Braitenberg, 1962; Vogt & Vogt, 1919) architecture, and which are heavily folded within the compressed cranial space. Advances in high-resolution MRI scanning have continued to improve our capacity to evaluate the anatomical and areal composition of the cortex in its compacted form, including using dMRI in living subjects (Wang et al., 2021). However, despite rapid progress in biophysical modelling (Jelescu et al., 2022; Zhang et al., 2012) of the dMRI signal, many open questions remain about how the detailed anatomical features of the cortex influence the local diffusion parameters. Gathering this information will help the development of biophysical models that inform basic neuroscience research, and improve the diagnosis and tracking of brain disease (Andica et al., 2020; Liu et al., 2018; McKavanagh et al., 2019; Torso et al., 2023; Weston et al., 2020).

One important feature of interest is the myelin composition of the cortex, the density and laminar distribution of which varies between areas (Bock et al., 2011; Braitenberg, 1962; Glasser et al., 2014). Myelin is a composite of lipids and proteins that coats a subset of cortical axons (Kandel et al., 2013; Morell & Quarles, 1999), primarily to facilitate the passive conduction of action potentials, and changes in cortical myelin content play important roles in development, plasticity and disease (Kandel et al., 2013; Morell & Quarles, 1999). Because myelin presents a stronger barrier to the diffusion of water molecules than the cell membrane, high-resolution dMRI might potentially be used to map the myelin density at each location in the brain, though a reliable method for this has not yet been established.

At present, there are a number of non-diffusion MRI strategies to map myelin content in the cerebral cortex. These include the T1w signal (Bock et al., 2011; Glasser et al., 2014), myelin water fraction (Heath et al., 2018; Lazari & Lipp, 2021; Mancini et al., 2020) and the magnetization transfer ratio (**MTR**) (Grossman et al., 1994; Heath et al., 2018; Lazari & Lipp, 2021; Mancini et al., 2020). Magnetization transfer imaging estimates the density of protons bound to large macromolecules based on the frequency shift they undergo (Grossman et al., 1994; Wolff & Balaban, 1989). In a previous study, we found that the MTR signal was strongly correlated with myelin density in co- registered histological sections of the cerebral cortex (Reveley et al., 2022).

Parameters from diffusion tensor imaging (**DTI**), such as fractional anisotropy (**FA**), radial diffusivity (**RD**) and axial diffusivity (**AD**) have also been used for estimating myelin content. However, studies have found varying levels of correlation between these parameters and histological myelin levels. In a previous study (Reveley et al., 2022), we found that in general DTI parameters correlated poorly with myelin density, although radial diffusivity showed a consistent relationship with a measure of axon organization derived from myelin histology.

The DTI model assumes that the distribution of diffusing molecule displacements is Gaussian. However, the physical barriers imposed by some cellular features, most notably myelin, may restrict diffusion and thus create differently shaped distributions. A wide variety of non-Gaussian modelling approaches are possible for dMRI data acquired with multiple b-values. The most conceptually straightforward of these is diffusion kurtosis imaging (Jensen et al., 2005), which offers scalar parameter maps of mean, radial and axial kurtoses, summarizing deviation from a Gaussian displacement distribution. Some studies have reported a link between these parameter maps and tissue myelin levels in murine white matter (Falangola et al., 2014; Guglielmetti et al., 2016; Kelm et al., 2016). In addition, the kurtosis measures are also frequently linked to “tissue complexity” (Wu & Cheung, 2010) and have been employed to characterize a range of conditions including astrogliosis, glioma, multiple sclerosis, stroke and neurodegeneration (Zhuo & Gullapalli, 2020)

Here, we investigated the relationship of one such diffusion statistic, the mean diffusion kurtosis (**MK**), to the myelin distribution within the cortical gray matter. We acquired multi-shelled dMRI scans of the fixed brains of two common marmosets (Callithrix jacchus) at spatial resolutions of 80-150µm. Tissue myelin levels were estimated with MTR scans in the same brains at 75-80µm. Subsequently we sectioned, stained for myelin, and co-registered the tissue of one of the brains, allowing for a pointwise comparison of the dMRI parameter maps and tissue myelin levels across the grey matter. We found that MK, and additionally parameter maps from the MAP-MRI (Özarslan et al., 2013) and NODDI (Zhang et al., 2012) models, closely tracked the myelin content in the cerebral cortex, as assessed both by direct histological comparison and comparison to precisely registered MTR data, used as a surrogate for myelin.

## Results

Our principal goal was to determine which diffusion MRI (**dMRI**) parameters most closely track the density of myelin in the gray matter tissue of the cerebral cortex. To this end, we analyzed high resolution, ex-vivo dMRI data from the brains of two marmoset monkeys. We then compared this dMRI data to the myelin content of the tissue, first as assessed by a non-diffusion MRI sequence, and second as assessed by myelin-stained histology gathered from one brain, and carefully co-registered to the MRI (Reveley et al., 2022). We employed the magnetization transfer ratio (**MTR**) as our non-diffusion MRI measure, since this sequence is known to be sensitive to large macromolecules such as those comprising myelin (Grossman et al., 1994; Heath et al., 2018; Henkelman et al., 2001). In previous work we showed that MTR intensity closely matched the fine-scale distribution of myelin in the cortical sheet, using histology data gathered from the scanned tissue (Reveley et al., 2022).

### MTR tracks histological myelin in the cortex

We began by re-establishing that MTR is a valid surrogate measure for myelin intensity in the cerebral cortex. To do this, we directly compared the spatial distribution of MTR intensity to the non-linearly registered myelin-stained sections obtained from the scanned brain. This comparison is shown for one segment of the cerebral cortex in **Figure 1 A**, where the myelin-stained histological section (top) has been resampled to match the resolution of the MTR scan (bottom) at 75µm per pixel. Myelin was most intense in deep cortical layers, declining towards the pial surface in the same way in both histology and MTR data. In general, the different modalities showed strong agreement in both the laminar and tangential aspects of cortical variation, for example in the position of the middle temporal (MT) area, which is known for its high myelin content. **Figure 1B** shows a pixel-by-pixel comparison of the two modalities, demonstrating a strong correlation (**ρ=0.94**). We found that each of the registered myelin-stained sections showed a strong correlation between MTR and histological data (**SI Figure 1**), indicating that the high-resolution MTR acquisition from fixed brains is a reliable proxy for cortical myelin distribution.

**Figure 1.**
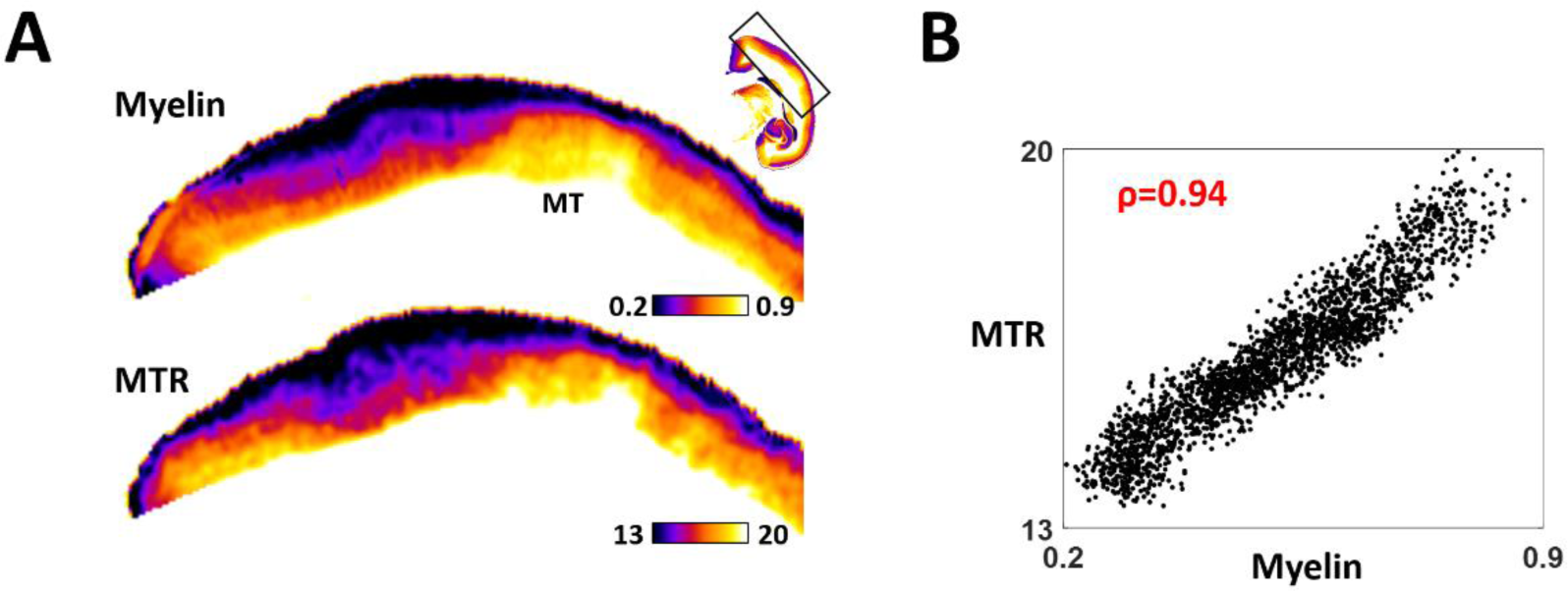
Histological myelin is strongly correlated with Magnetization Transfer Ratio (MTR) MRI. **A:** MTR and myelin histology (Gallyas stain) non-linearly registered at 75µm resolution, showing area MT. Myeloarchitectural features (top row) show strong congruence with MTR intensity (bottom row) **B:** Pixelwise scatter plot of the data shown in A) shows a very strong correlation and linear relationship (ρ=0.94). Correlations for all seven myelin-stained sections are shown in **SI Figure 1**

With this tool in hand, we focused on our target question of how diffusion signals in the cortex might reflect myelin content as measured by either MTR or histology. We previously demonstrated that one common diffusion measure, fractional anisotropy (FA) of the Gaussian diffusion tensor, was a poor predictor of myelin content (Reveley et al., 2022). Our present hypothesis is that non-Gaussian diffusion - specifically restricted diffusion - may track myelin content in the cortex.

### Mean diffusion kurtosis tracks myelin levels

The restriction of water diffusion due to periaxonal myelin sheets is shown in two dimensions in **Figure 2**. Briefly, during free diffusion, water molecules exhibit random walks (**Figure 2A**), such that their mean squared displacement increases linearly with time, which is reflected in a Gaussian distribution of molecule displacements at time T. In biological tissue, diffusion is often hindered by tissue features. Although hindering diffusion may reduce the mean squared distance travelled by molecules in a particular direction, the molecule displacements remain close to Gaussian, and this can be effectively modelled by the diffusion tensor. However, when a strong diffusion barrier such as myelin is present, molecule displacement within the axoplasm is severely restricted (Schneider & Wheeler-Kingshott, 2014). In this case, the maximum displacement is bounded by axon size (**Figure 2B**) (Novikov, 2021). In restricted diffusion, the mean squared displacement of the restricted molecules over time becomes sublinear, and the displacement distribution of all molecules at time T becomes kurtotic. The mean diffusion kurtosis (**MK**) stemming from restricted diffusion can be estimated across the tissue using multi-shell dMRI acquisition (Jensen et al., 2005) (see **Methods**).

**Figure 2.**
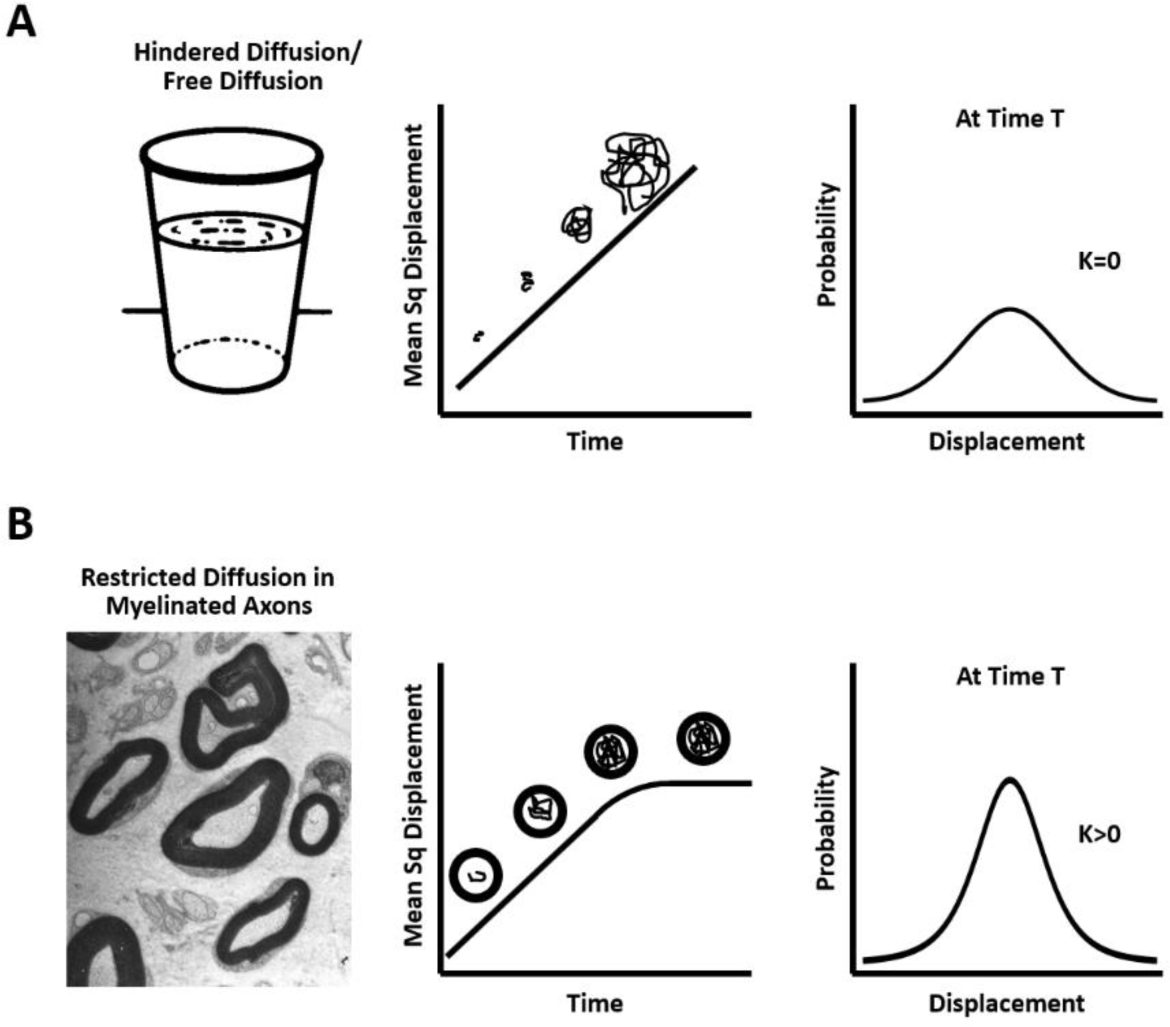
Idealized properties of free/hindered diffusion and restricted diffusion with respect to myelin. **A:** In free and hindered diffusion, the mean squared molecule displacement is linear with respect to time. At a given time T, the probability distribution of molecule displacements is Gaussian, which can be captured by diffusion tensor imaging. **B:** In the presence of myelinated axons, the maximum distance a molecule can travel is bounded by the size of the axon, and the displacement distribution at a given time becomes kurtotic. Figure modelled after *(Schneider & Wheeler-Kingshott*, *2014)*, image from *(Reina et al., 2015)*

We compared the MK mapped across the whole cerebral cortex with the myelin content as estimated by the MTR scan. Since both maps are MRI measures from the same ex-vivo sample, the spatial registration was very precise. Importantly, the MK and MTR measures capture fundamentally different physical processes, related to the water diffusion profile and off-resonance spin exchange coming from large molecules (Henkelman et al., 2001) , respectively. Nonetheless, we found the two measures exhibited strong similarity in the monkey cortex.

Figure 3A and B illustrate this similarity in representative coronal, sagittal and axial slices at 80 mm resolution. Within these sections, the close correspondence in both the laminar and tangential features is evident. In the laminar direction, the MK and MTR measures both exhibited the expected myelin gradient, showing high values in the lower layers near the white matter and decreasing values toward the pia. In the tangential direction, the changes across cortical areas were also similar. For example, the elevated MTR in MT (middle temporal cortex) and M1 (primary motor cortex) were reflected by elevated MK in the same areas, due to the putative restricted diffusion within the relatively more numerous axonal myelin sheaths in these cortical regions.

**Figure 3.**
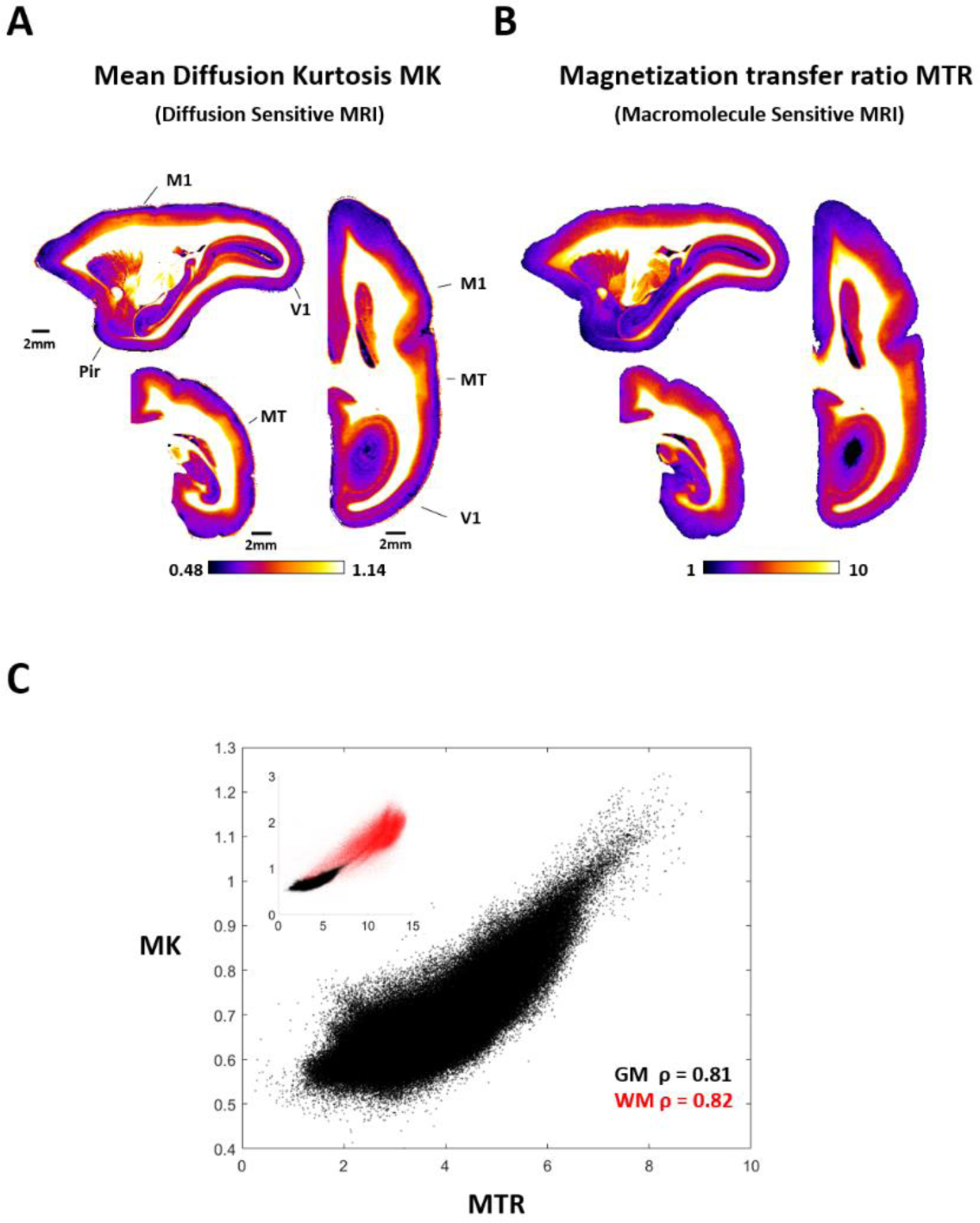
Mean diffusion kurtosis and the magnetization transfer ratio have a similar spatial profile: **A:** The distribution of mean kurtosis at 80 µm matches the expected myelin distribution in highly myelinated (Primary motor **M1**, middle temporal **MT**), moderately myelinated (Primary vision **V1**) and lightly myelinated (Piriform cortex, **Pir**) cortical regions. **B:** The distribution of MTR is visually similar to MK, especially in high myelin areas. **C:** Intensity of MK and MK voxels plotted against each other in gray matter (black, ρ=0.81) and white matter (red, ρ=0.82). Inset shows gray matter and white matter together.

We quantified the pixelwise relationship between MK and MTR (Figure 3 **C**), which revealed a strong correlation in the gray matter (**ρ=0.81**, black Figure 3 **C**). The mean kurtosis measure therefore appeared sensitive to variation in the vertical and horizontal cortical myelin distribution, in a similar way to the MTR. Interestingly, when this analysis was extended to the white matter, with its higher overall myelin content due to dense composition of myelinated axons, the correlation remained strong (**ρ=0.82** red inset Figure 3 **C**). White matter pixels with higher MTR values, reflecting a higher spin transfer from large molecules, also exhibited higher MK, reflecting more restricted water diffusion. This robust agreement between diffusion and large-molecule MRI measures in the precisely registered maps of the same brain suggests that mean diffusion kurtosis is a reliable and quantitative marker for myelin content. At the same time, the strength of the correlation was slightly weaker than that between myelin histology and MTR, shown in Figure 1 **B**. To investigate this discrepancy further, we examined the regional variation in the relationship between MK and MTR.

We divided the cortex into anatomical regions based on the areal definitions of the Paxinos atlas (Paxinos et al., 2013) and marmoset brain mapping project (Liu et al., 2018, 2020), using the registered maps provided by a previous study(Liu et al., 2018, 2020). We found that the correlation coefficient between MK and MTR varied across cortical regions (Figure 4 **A**). In contrast to a surface-based analysis of values averaged through the cortical depth, our voxel-based methods at ultra-high resolution captured rich laminar detail of the cortex (Figure 4 **B**). Nevertheless, the variation in regional correlation coefficients was related to each region’s total myelin content. For example, in cortical regions where the mean MTR value was high, and which were known to have high myelin content, for example M1 or MT (Figure 4 **B**) correlation levels approached 1, whereas regions with low MTR, such as the piriform cortex (Pir) exhibited lower correlations, closer to 0.6. The lower correlations may be linked to a decline in contrast relative to noise where myelin content is lower, rather than a failure of either measure to track myelin content (**SI** Figure 2).

**Figure 4.**
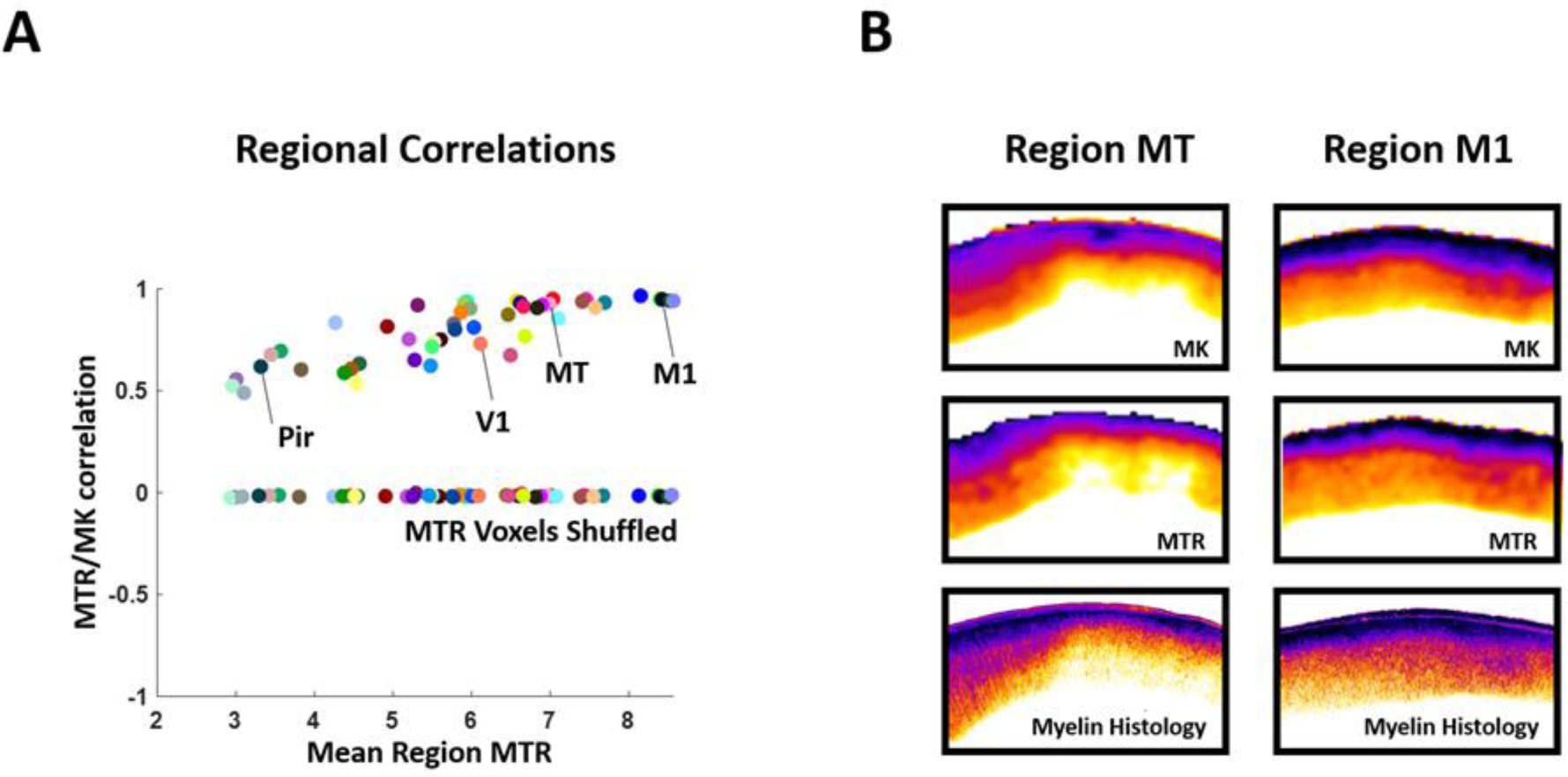
Correlations of MK and MTR at 80 µm resolution for each cortical region. **A**: Pearson correlation coefficients between MK and MTR for each marmoset cortical region as delineated in Liu et al *(Liu et al., 2018, 2019)*, plotted against the mean MTR value for each region. Higher correlation coefficients were observed for regions with higher mean MTR values. A simple shuffle control was implemented by randomly shuffling the voxel order of the MTR data. See color key in SI Figure 2. **B**: MTR, MK and myelin histology examples from two cortical regions M1 (Primary motor) and MT (middle temporal).

### Comparison with related diffusion measures

We next expanded our investigation of dMRI and cortical myelin to include two further popular diffusion models, namely the non-gaussian (**NG**) component of the mean apparent propagator (MAP- MRI) MRI model (Özarslan et al., 2013) and the neurite density index (**NDI**) calculated as part of neurite orientation dispersion and density imaging (**NODDI**) (Zhang et al., 2012). The NG parameter computed

from MAP-MRI is conceptually similar to mean kurtosis. However, NDI is not a measure of non-Gaussian diffusion per se, but is instead part of NODDI, a biophysically motivated tissue model of Gaussian diffusion that seeks to estimate the variation in neurite density (Zhang et al., 2012). In NODDI, contributing gray matter neurites can potentially include both myelinated and unmyelinated axons, as well as dendrites, all of which are modelled as sticks with negligible radial diffusivity (Zhang et al., 2012).

We found that all three of the dMRI parameter maps we considered (MK, NG and NDI) resembled the MTR and one another in the cortical gray matter, but differed substantially from the FA maps of the diffusion tensor (Figure 5 **A**). The correlation between the three dMRI measures of interest was extremely high, with the correlation between MK and NDI at 0.86 and between MK and NG at 0.92. The correlations with each of these parameters to the MTR maps were also high, albeit with the correlation of NDI to MTR taking a slightly lower value (0.73) than those of MK and NG (0.81 and 0.8 respectively). By contrast, the relationship of FA to the three putatively myelin sensitive dMRI measures, and to MTR, were all much lower, between 0.26 and 0.3 (Figure 5 **B**). When we examined the spatial relationships of dMRI parameters to MTR, we found that MK and NG exhibited a nearly identical, roughly linear voxel wise relationship to MTR, and NODDI showed a slightly different but still similar relationship. The relationship of FA to MTR, however, was much weaker (Figure 5 **C**).

**Figure 5.**
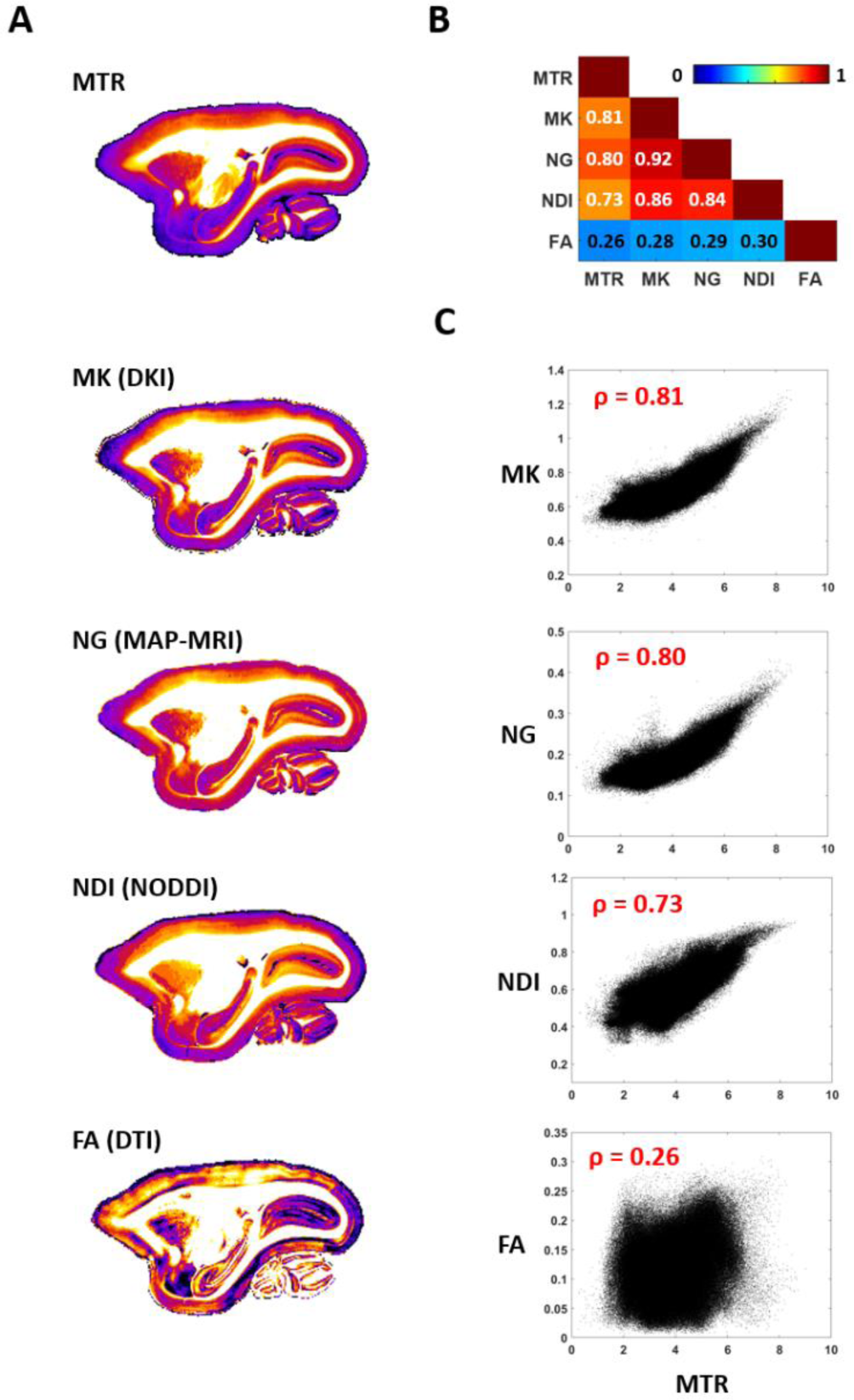
The parameter maps NG from MAP-MRI and NDI from NODDI show a similar relationship to MTR as mean kurtosis (MK) **A:** Example slices of MTR, MK, NG, NDI and FA from the marmoset. The first 4 are very similar in the gray matter, while the FA shows a different cortical profile. **B:** Pearson correlation coefficients between all parameter maps are strong, except those involving FA **C:** When plotted against MTR, NG and MK both show a very similar linear relationship. The relationship of NDI to MTR is slightly different, but still qualitatively similar. In contrast, the correlation between FA and MTR is poor and the values appear randomly distributed. The analysis included only gray matter.

### Direct comparison to histological myelin content

Having established the close relationship between MTR and the diffusion parameters MK, NG and NDI over the whole brain, we next focused on a fine-grained comparison of the dMRI parameters with myelin histology obtained from the same brain. We compared 7 non-linearly registered sections of myelin histology to matching dMRI and MTR MRI. In the example scatter plots from one section shown in Figure 6 **A** the MRI parameters of interest all showed a clear linear relationship with histological myelin, with MTR, MK, and NG showing Pearson correlations of 0.85, 0.82 and 0.84 respectively. NDI showed a slightly weaker correlation of 0.75. FA showed a lower overall correlation and a more complex, piecewise relationship to myelin density, which we previously reported (Reveley et al., 2022).

**Figure 6.**
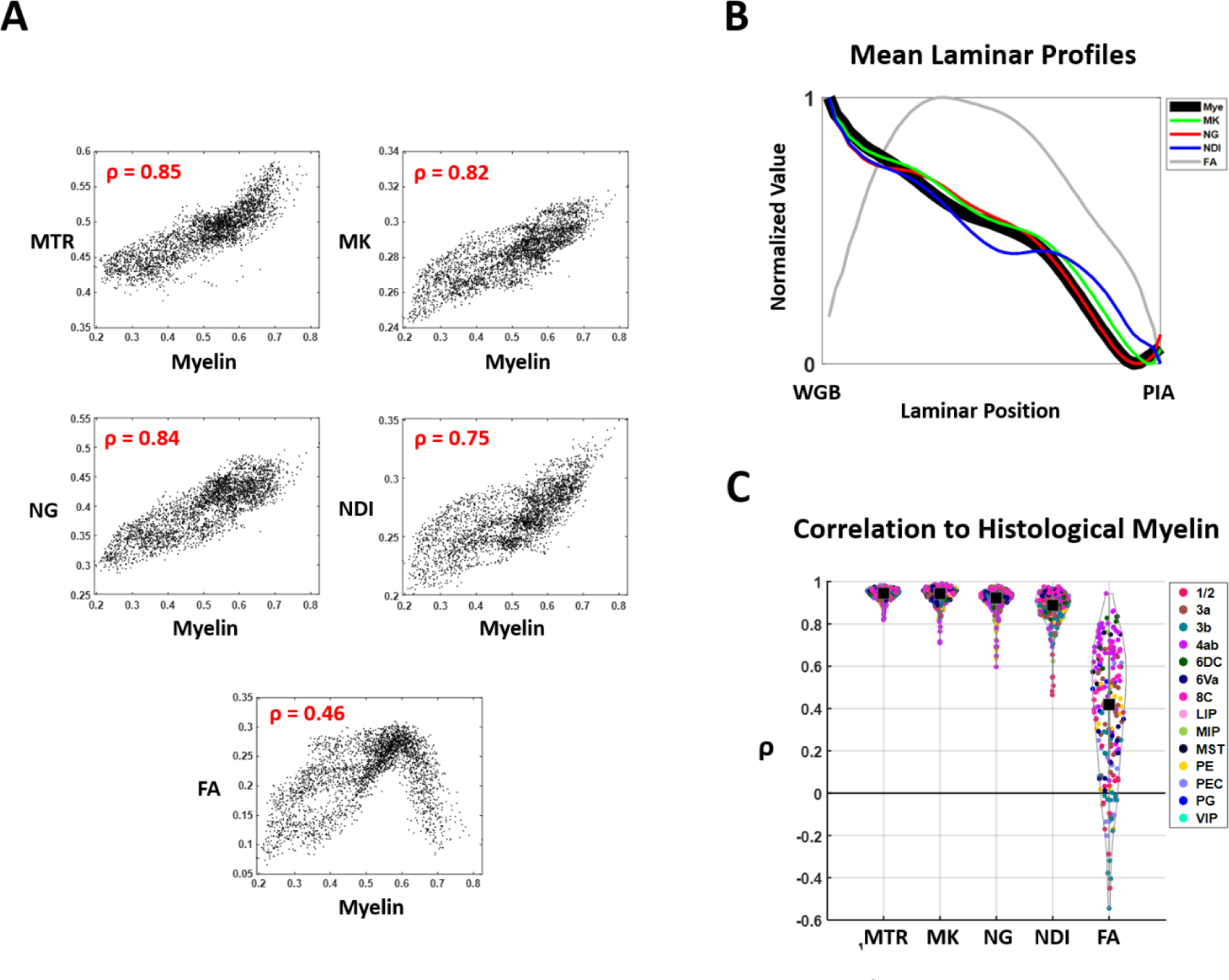
The parameter maps MTR, NG, MK and NDI show a similar relationship to histological myelin. **A:** Pixelwise scatter plots of MTR, NG, MK, NDI and FA plotted against histological myelin in the gray matter of one histological section. **B:** The mean laminar profiles (see Methods) of histological myelin, MK, NG, NDI and FA. All the parameters follow a similar monotonic decline to myelin, except FA C: MTR, MK, NG, NDI and FA cortical parcel correlations to histological myelin (see **Methods**).

Continuing to investigate the histological basis of the dMRI parameters, we next examined their laminar variation, their variation by cortical area, and how both of these tracked histological myelin content. To assess the similarity in laminar variation, we sampled the cortex along columnar lines derived from 2D tractography, and resampled the resulting data into a matrix formulation (see **Methods**). This allowed us to calculate and compare the mean laminar profile of the various MRI measures to the co-registered myelin histology (Figure 6 **B**). We found that NG and MK exhibited a laminar profile that was nearly identical to that of myelin intensity, declining monotonically from deep to superficial layers. However, the laminar profile of FA differed substantially from that of myelin, and from the other diffusion parameters, with minima at the white/gray and pial boundaries and a peak in middle cortical layers (Reveley et al., 2022).

To provide a more granular analysis, we next sampled the matrix representation of the cortex into columnar parcels (Reveley et al., 2022) running vertically from the white matter to the pia and with a horizontal width of about 300 µm (Figure 6C). Within each of these parcels, we measured the

correlation of the MRI parameters with histological myelin. This analysis showed a high spatial correlation of voxels within most columnar parcels, with the highest median values observed for MTR (0.95) and MK (0.94), slightly lower values for NG (0.92) and NDI (0.89), but much lower values observed for FA (0.42).

Together, these analyses reveal that the diffusion-based measures - obtained either by modeling the spatial distribution of diffusion (MK, NG) or by biophysical forward modeling of the neurite distribution (NDI) - were all good measures of the cortical myelin distribution. This result is especially clear when the higher order parameters are contrasted to FA, which does not correlate with myelin levels in a straightforward way.

### Comparison to radial diffusivity

The above results show that several higher order models obtained from multi-shelled diffusion data closely track myelin content in the cortex. Although FA in the cortex does not track myelin density, radial diffusivity (**RD**), another measure derived from the diffusion tensor, has been routinely used for this purpose. In some studies, RD has been shown to correlate with histological measures of myelin content and has been used as a marker for assessing myelin integrity and alterations in white matter microstructure (Lazari & Lipp, 2021; Mancini et al., 2020). Previously, we found that RD tended to track myelin organization rather than myelin intensity (Reveley et al., 2022). We next examined the relationships between RD and myelin, using the same approach as above (Figure 6).

We found that although RD does show a correlation to myelin intensity, this correlation is notably weaker than MK’s, exhibiting a much wider spread of correlation coefficients between columnar parcels of cortex (Figure 7A). The laminar profile of RD also diverged significantly from that of myelin histology (Figure 7B**, C**). We found that while the spatial pattern of RD was very sensitive to the directional content (Reveley et al., 2022) (**SI** Figure 3, **SI Text 1**) of the myelinated axons as well as their density, it did not serve as a good quantitative measure of myelin content in the same way as mean kurtosis.

**Figure 7.**
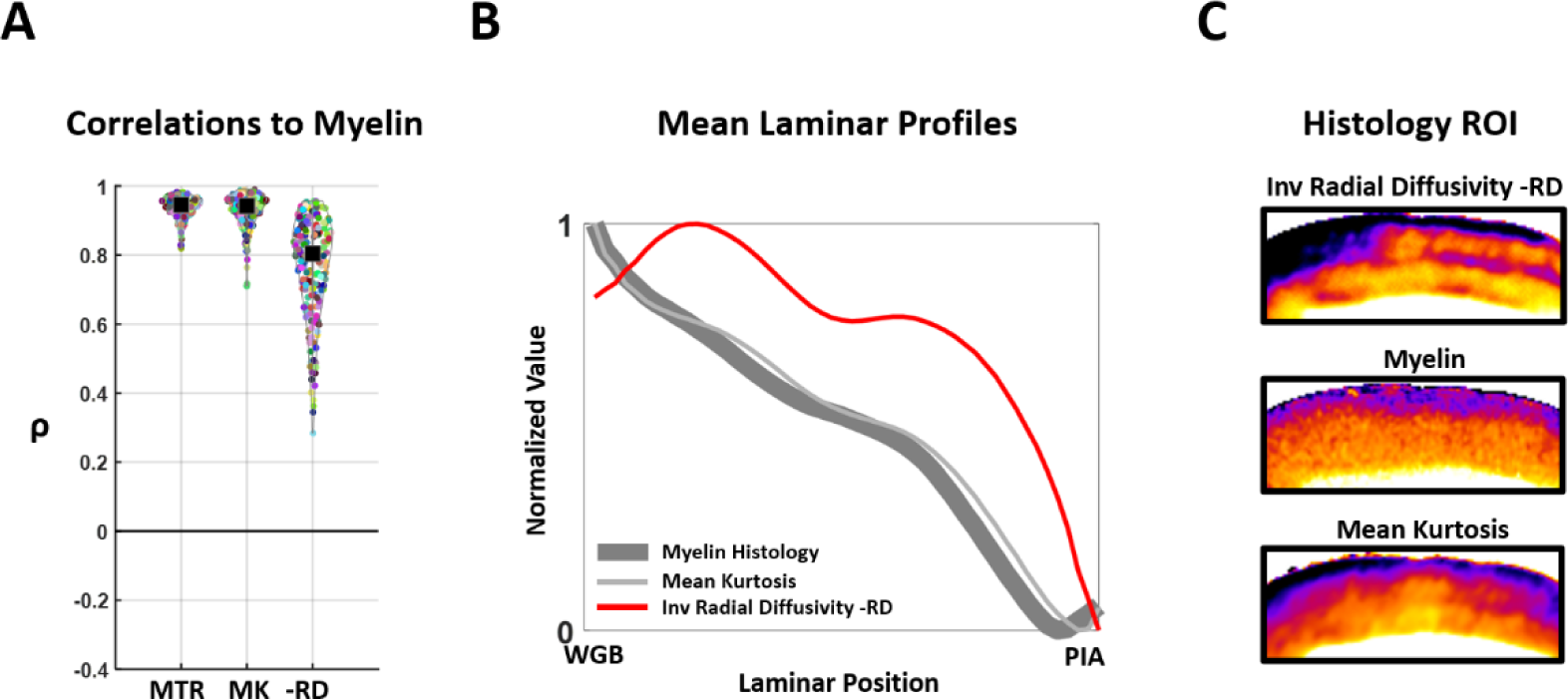
Differences in the relationships of RD and MK to histological myelin. **A:** Cortical parcel correlations (see **Methods**) of MTR, MK and inverse radial diffusivity (-RD) to histological myelin. -RD shows lower mean correlation, and a greater spread of values **B:** The mean laminar profiles (see **Methods**) of Histological myelin, MK and -RD C: Representative ROI of histological myelin, MK and - RD.

### Mean diffusion kurtosis in area V1

There is wide regional variation in cytoarchitecture, myeloarchitecture and neurite structure over the expanse of the cortical sheet. We were interested whether MK could assess the laminar structure of the primary visual cortex (**V1**), a well-studied cortical area with a unique laminar patterning of cell density and myelin (Balaram & Kaas, 2014; Casagrande & Kaas, 1994). Figure 8 demonstrates the capacity of MTR, MK, NDI and FA to reveal aspects of V1’s laminar architecture in the postmortem sample. Importantly, the laminae of highest intensity did not match among all the registered MRI measures. MTR, MK and NDI matched well and showed their highest levels in layer 4B, the so called Gennari band, reflecting its known high myelin content (Balaram & Kaas, 2014; Casagrande & Kaas, 1994) (Figure 8A).

**Figure 8.**
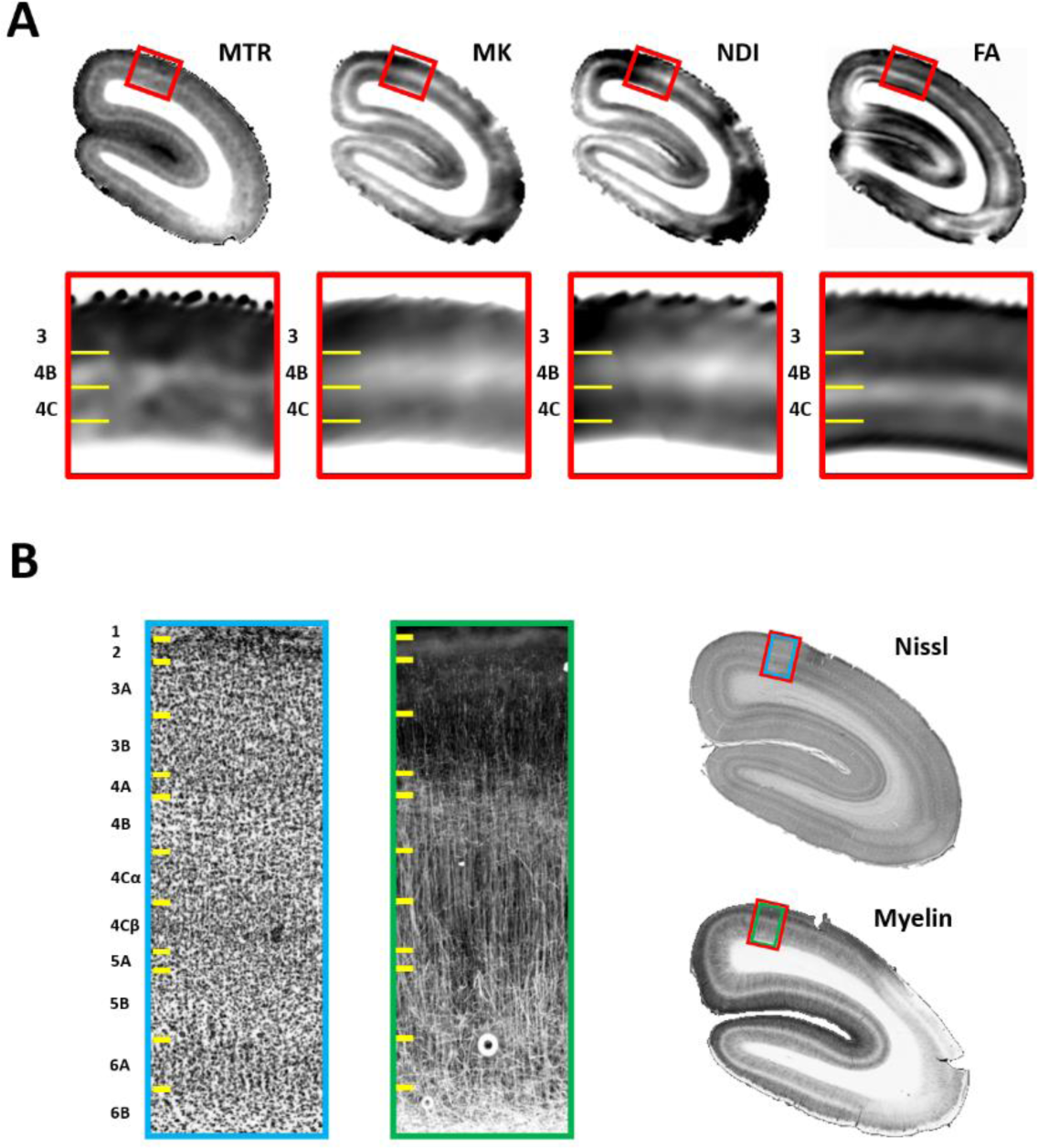
MTR, MK, NDI reveal the same laminar properties of myelin in primary visual cortex, while FA appears to reflect non-myelinated components. **A:** MTR, MK, NDI and FA parameter maps expose different cortical layers and sublayers. In particular, the high myelin Gennari band of layer 4B is exposed as a high intensity band of MTR, MK and NDI, but a low intensity band of FA. In contrast, Layer 4C is shown as a low intensity band of MTR, MK and NDI but a high intensity band of FA. **B:** Nissl and myelin histology from the same subject matched to the MRI shown in A. See text for discussion.

The abundant myelinated axons in 4B are visible in the corresponding histology to have a matted structure (Figure 8 **B**) and this lack of spatial coherence explains the low FA in this layer, and also in the deepest layer of cortex adjacent to the white matter. By contrast, layer 4C was revealed by a high intensity band of FA, which likely reflects an abundance of vertically oriented, non-myelinated neurites in this layer (Figure 8A**, right**) (Reveley et al., 2022). Additionally, we found that radial diffusivity was relatively lower in layer 4C where the myelinated axons are coherently organized, than in 4B where they are matted, despite the higher myelin content of the Gennari band in 4B (**SI** Figure 4**, SI Text 2**). This likely reflects the sensitivity of RD to the organization of the myelinated fibers as shown in Figure 7 **B**. Together, these results indicate that models based on multi-shelled diffusion acquisitions can distinguish and quantify the diffusion profiles of myelinated and non-myelinated neurites in the gray matter at a laminar level.

## Discussion

In this study, we assessed the relationship between cortical myelin content and higher order diffusion MRI model parameters based on multi-shelled acquisitions. We analyzed ultra-high-resolution dMRI and MTR scans of whole primate brains, as well as co-registered histological data covering large areas of the frontal and parietal lobes. We examined the differences between cortical regions, as well as the laminar properties of the parameters within those regions. Our guiding hypothesis was that water restricted to the intra-axonal space by the myelin sheath, would be reflected in a non-Gaussian molecule displacement profile, so we initially focused our investigation on mean kurtosis (MK). MK strongly correlated with MTR - a separate MRI contrast unrelated to diffusion, but closely linked to myelin levels - globally over the whole cortical sheet. Regions with the highest myelin content, and therefore the highest signal-to-noise ratio, exhibited the strongest correlation with MK. Broadly similar results were obtained with the non-Gaussian (NG) parameter map obtained from MAP-MRI (Özarslan et al., 2013), and neurite density index (NDI) parameter map obtained from the NODDI biophysical model (Zhang et al., 2012). Each of these high order diffusion measures closely tracked myelin, as well as one another. Specifically, they matched the monotonic decay of histological myelin levels from the white gray boundary to the pial surface, as well as the tangential variation across the cortical sheet. In the primary visual cortex, the MTR and MK measures captured the known distribution of myelin fibers in layer 4B. Of the three higher order diffusion parameters we tested, NDI was unlike MK and NG since it does not explicitly reflect non-Gaussian diffusion processes. Potentially, both myelinated and unmyelinated neurites might drive the NDI contrast (Zhang et al., 2012), but our analysis shows that NDI behaves similarly to MK, a measure of non-Gaussianity, in the environment of fixed primate gray matter.

The behavior of MK, NG and NDI may be contrasted with the behavior of the Gaussian diffusion tensor in gray matter. Radial diffusivity (RD) exhibited sensitivity to the organization of the myelinated fibers as well as their density, rather than being attuned to myelin density alone. Fractional Anisotropy (FA) showed only weak correspondence to myelin parameters, and in previous work we showed that FA is strongly driven by non-myelinated tissue components. Employing both the diffusion tensor and higher order diffusion terms may offer a way to separate out the specific influences of myelinated and non-myelinated tissue components on the diffusion signal. Moreover, we found that MTR and MK were both myelin sensitive despite obtaining their contrasts via different mechanisms. Thus, although they have correlated values in healthy tissue, MK and MTR might diverge for pathological cases in specific ways that have diagnostic value (Edwards et al., 2018; Kamiya et al., 2020).

### Relation to previous findings

Our results cohere with, and draw together several lines of converging evidence (Fukutomi et al., 2018; Guglielmetti et al., 2016; Jelescu et al., 2016; Kelm et al., 2016; Yoshida et al., 2013) at different spatial scales, to suggest that mean kurtosis has both sensitivity and specificity to myelin levels in healthy gray matter.

Mean kurtosis values have previously been shown to track changing myelination levels *in vivo*. For example, Guglielmetti et al (Guglielmetti et al., 2016) studied dMRI and immunohistochemical material from mice which had been treated with cuprizone, a drug that reversibly induces demyelination through toxicity to oligodendrocytes. The authors found that mean kurtosis decreased in the gray matter of motor and somatosensory cortex of the mouse following cuprizone administration, and regained their values on recovery from the treatment. This was true except in areas of cortex where the histology showed remyelination to be incomplete. Likewise, in a developmental study, Praet et al (Praet et al., 2018) assessed the correlation of mean kurtosis based on the optical density of co-registered myelin basic protein immunohistochemistry. They found that as myelin levels in the motor cortex increased during development, cortical MK tended to increase and FA tended to decrease. These rodent studies emphasized longitudinal changes in experimental cohorts, rather than detail within the cortex of particular subjects.

A few studies have examined the distribution of higher order diffusion metrics in groups of human subjects at clinical resolutions over the whole brain. Bester et al (Bester et al., 2015) examined diffusion kurtosis in a cohort of subjects suffering from multiple sclerosis (MS), a demyelinating disease. They found that MK was significantly lower in the gray matter of MS patients compared to controls, and that MK was inversely correlated with executive function where poor function presumably reflects cortical demyelination. Fukutomi et al (Fukutomi et al., 2018) found that NDI was strongly correlated to myelin density as estimated by the T1w/T2w ratio, in a group based cortical surface based study of 505 healthy subjects from the human connectome project. These studies both suggest that the present findings in an ex-vivo primate model are strongly relevant to human cortical tissue scanned in-vivo, at much lower spatial resolutions.

Zhu et al (Zhu et al., 2021) compared MK in a cohort of six postmortem macaque brains at lower spatial resolutions of around 600 µm in plane, gathered in 2mm slices, to publicly available neurofilament (SMI-32) immunohistochemistry from a different animal. They found qualitative correspondence between the two measures over the cortex as a whole, however they were not able to examine the histology/dMRI relationship at a fine spatial scale in the same brains. SMI-32 is reactive in a subset of large pyramidal cells in the cortical gray matter, and is closely associated with myelination (Kirkcaldie et al., 2002), because those same cells give rise to the largest myelinated axons in cortex. The broad correspondence Zhu et al noted at comparatively low spatial resolution may be a proxy for average cortical myelin levels, rather than indicating a direct link between MK and the neurofilaments themselves. More detailed work is required to address this question.

A handful of studies (Lazari & Lipp, 2021) have examined the relationship of diffusion kurtosis to the detailed properties of myelinated axons in murine white matter using electron microscopy. Kelm et al (Kelm et al., 2016) studied the relation of myelin to diffusion kurtosis in an ex-vivo cohort of knockout and control mice with varying degrees of hypomyelination. They found that MK correlated to the fraction of myelin as estimated from the electron microscopy data in four white matter ROI across their subject pool. Jelescu et al (Jelescu et al., 2016) also found that MK exhibited sensitivity to cuprizone induced demyelination in an analysis of electron microscopy in the mouse corpus callosum.

### Caveats and limitations of the present study

Connecting MRI data to underlying tissue parameters requires certain assumptions, so it is important to identify where such assumptions might break down. Importantly, while the focus of the present study was on myelin, both MTR and MK may also have sensitivity to other tissue properties. For example, MTR is known to vary with water content, as brought about by edema, inflammation, and other immune responses (Heath et al., 2018; Vavasour et al., 2011). Likewise, beyond marking axonal myelination, MK varies with other factors that increase tissue “complexity”, such as gliosis or gliomas (Wu & Cheung, 2010). These caveats may be particularly important in the translation of the *ex vivo* data presented here to the living brain. Isolating any myelin-unique component of MRI signals is particularly difficult in the cortical gray matter, because multiple tissue parameters correlate spatially with one another. For example, in the cortex, the largest myelinated and non-myelinated neurites both have a strong vertical orientation bias and are spatially enmeshed (Buxhoeveden & Casanova, 2002; Peters et al., 1997; Peters & Sethares, 1996). Nonetheless, our demonstration of a laminar gradient of MK in the cortex does suggest that myelin is the determining influence, since this is a signature of cortical myelin levels (Braitenberg, 1962), whereas the density of unmyelinated neurites is typically high in superficial cortical layers, near the pia (Peters, 2010; Peters et al., 1997).

Diffusion parameters are sensitive to the state of the tissue. In the present study we exclusively addressed fixed, *ex vivo* tissue scanned at diffusion times in the range 10-20ms (see **Methods**). Acquisition in living tissue might give contrasting results, since both diffusivities and higher order diffusion measures differ in living tissue. We note that Fukutomi et al found good agreement between NODDI NDI and myelin as estimated by the T1w/T2w ratio over the whole human brain in-vivo (Fukutomi et al., 2018), so far the ex-vivo and in-vivo findings are supportive to each other. Work to assess these issues and develop sophisticated, empirically grounded models of complex diffusion processes in gray matter is ongoing (Jelescu et al., 2022; Lee et al., 2020).

## Methods

### Methods summary

The postmortem brains of two adult marmosets “case M” and “case P”, both used in previous studies (Liu et al., 2018, 2020; Reveley et al., 2022), were employed in this work. Both animals were scanned for multi-shelled diffusion MRI (dMRI) and for Magnetization Transfer Ratio (MTR) MRI on the same 7T Bruker preclinical scanner with a 30cm bore and 450mT/m gradients. Case M was scanned and then carefully co-registered to the Paxinos atlas of the marmoset cerebral cortex as part of the marmoset brain mapping project (Liu et al., 2018, 2020). The dMRI was obtained at a resolution of 80µm isotropic in a whole brain scan lasting 15 days, for that project (Liu et al., 2020). The left hemisphere of “case P” was scanned specifically for the present study at a resolution of 150 µm isotropic. As part of a previous study (Reveley et al., 2022), case P was sectioned for Gallyas myelin histology, which was then non-linearly registered to the dMRI data. All the procedures and provision of materials for this study were in full compliance with the Guidelines for the Care and Use of Laboratory Animals by National Institute of Health and approved by the Animal Care and Use Committee of the National Institute of Mental Health.

### Ex vivo high-resolution MRI Acquisition

#### Case M

Detailed scan information can be found in Liu et al 2020 (Liu et al., 2020). Briefly, the brain of case M (male common marmoset , 4.5 years old “Case M”) was soaked with 0.2% gadolinium (1 mmol ml–1 Gadavist, Bayer) in 1× PBS for 2 weeks and then with 0.2% gadolinium in pure water for 1 day. Case M was then scanned using a coil specially tailored for ex-vivo scanning of marmoset brains. 204 dMRI volumes were acquired at 80 µm isotropic resolution in four shells (8 b = 0, 6 b = 30, 64 b = 2,400 and 126 b = 4,800). The pulse width for the diffusion weighting gradient was 6.4 ms and pulse separation is 14 ms, for a diffusion time of approximately 14 ms. The scan time was 15 days.

MTR data was acquired was acquired for case M, also at 80 µm isotropic resolution using a 3D FLASH sequence. Five MTR scans were acquired, each comprising two volumes with (Msat) and without (Moff) an offset magnetization transfer (±2,000 Hz off-resonance, Gaussian-shaped), which were averaged before calculating the MTR. The MTR value is calculated as 100(1-Msat/Moff). All five MTR scans were then averaged. The total acquisition time was 50 hours.

#### Case P

Before MRI scanning, the formalin-fixed marmoset brain of case P was soaked with 0.15% gadopentetate dimeglumine (Magnevist, Bayer Leverkusen, Germany) for 3 weeks to reduce the T1 relaxation time as described in a previous study(Liu et al., 2018; Reveley et al., 2022). In order to improve SNR, brain P was cut through the midline. Only the left hemisphere was used in the present study. The left hemisphere was scanned with a 25-mm birdcage volume coil (Bruker). The imaging parameters were: TR = 470 ms, TE = 34 ms, flip angle = 90°, FOV = 38.4 x 24 x 21 mm, matrix size = 256 x 160 x 140, resolution = 0.15 mm isotropic. The pulse width for the diffusion weighting gradient was 8 ms and pulse separation is 18 ms, for a diffusion time of approximately 18 ms. In order to keep the echo time short, 10 segments were used in the EPI acquisition. The DWI data were acquired using 2 averages, and the acquisition time for each 3D volume was about 22 minutes. In total, 142 DWI volumes were collected to sample the q-space on 5-shells. The 5 shells were defined by b-values: 419, 1677, 3773, 6708, and 10481 s/mm², obtained by setting the gradient magnitude at 80, 160, 240, 320, and 400 mT/m. 3, 21, 21, 37, and 58 volumes were collected in each shell respectively. Two volumes were collected at b=0.

The left hemisphere of case P underwent scanning for MTR at 75 µm as described in a previous study(Reveley et al., 2022). Briefly, The MTR images were collected with a 3D FLASH sequence, FOV = 34.8 x 20.4 x 20.1 mm, matrix size = 464 x 272 x 268. As with case M, the magnetization transfer was achieved in Msat scans by a Gaussian shaped pulse at +2000 Hz offset frequency in one scan and at - 2000 Hz in the other scan of a paired acquisition. This magnetization transfer pulse was turned off in Moff scans. The MTR value was calculated as 100(1-Msat/Moff). The number of averages was 5. The total MTR acquisition time for the left hemisphere of case P was about 60 hours.

### MRI processing

Here we describe MRI pre-processing steps and computation of diffusion tensor, kurtosis, NODDI and MAP-MRI parameters in each ex-vivo brain.

#### Case M

We used the publicly available 80 µm MTR and dMRI data from case M, which was previously preprocessed(Liu et al., 2020) using the DIFFPREP of TORTOISE (Pierpaoli et al., 2010). We removed the left hemisphere and closely cropped the right hemisphere using fslroi, part of FSL 6.0.5.2 (Woolrich et al., 2009), to save computation, since only the right hemisphere was aligned to the atlas labels. We calculated the mean, axial and radial kurtoses of the 80 µm data using dtifit part of FSL 6.0.5.2 (Woolrich et al., 2009). For the remaining parameter maps, we down sampled the diffusion data from 80 µm to 160 µm using the “fslmaths -subsamp2offc” command in FSL 6.0.5.2 (Woolrich et al., 2009), in order to save computation time. We computed the mean kurtosis a second time with “RobustDKIFitting” MATLAB script (Henriques et al., 2021) and compared this to the DTIFIT results, finding good agreement. We computed the NODDI model using AMICO (Daducci et al., 2015) version 1.5.4 with default diffusivity parameters and the “ex-vivo” parameter set to 1. We estimated the MAP model order 4 using TORTOISE (Pierpaoli et al., 2010) version 3.2.0, EstimateMAPMRI with default parameters (order 4).

#### Case P

For case P, we pre-processed the data using the DIFFPREP of TORTOISE (Pierpaoli et al., 2010). We registered MTR data to the histology slice plane and as described in a previous publication (Reveley et al., 2022), and performed a rigid body (6 parameter) registration of the dMRI data to the MTR using FSL 6.0.5.2 FLIRT (Woolrich et al., 2009), thus up sampling the dMRI from 150 to 75 µm with spline interpolation. We estimated the Mean Apparent Propagator (MAP) model at order 4 using TORTOISE (Pierpaoli et al., 2010) version 3.2.0, EstimateMAPMRI with default parameters (order 4). We computed the NODDI model using the NODDI (Zhang et al., 2012) toolbox 1.4, with parameters disoIdx 2.0E-9, diIdx 0.4E-9. We calculated the mean, axial and radial kurtoses of the 75 µm data using DTIFIT part of FSL 6.0.5.2 (Woolrich et al., 2009), and compared these results to those of the “RobustDKIFitting” MATLAB script (Henriques et al., 2021), finding good agreement.

### Anatomical mapping

#### Case M

The cortex of case M was carefully labelled and aligned to the Paxinos atlas in previous work (Liu et al., 2018, 2020). We removed the left hemisphere of the atlas MTR template and atlas labels since these were a mirror image of the right. We fine-tuned the alignment of the atlas labels and MRI data using a shift/translate only FLIRT transform with no interpolation. We used the medium granularity labels of the marmoset brain mapping labels (Liu et al., 2018) in this work. We manually trimmed the anatomical labels at the white gray boundary and pial boundaries to avoid partial volume effects and non-brain voxels. The white matter voxels were defined as part of the marmoset brain project white matter atlas (Liu et al., 2020). We down sampled each of the three label volumes released as part of the project to 160 µm using the “fslmaths -subsamp2offc” command in FSL 6.0.5.2 (Woolrich et al., 2009). We manually eroded each label sightly to avoid partial volume effects, and combined the three volumes into a single mask file.

#### Case P

For estimation of cytoarchitectural boundaries, we non-linearly registered the right hemisphere of case M MTR to the left hemisphere of case P MTR using ANTS 2.4.1 (Avants et al., 2011), as described in a previous study (Reveley et al., 2022). We then manually entered the coordinates of the region boundaries to the analysis code described below

### Histology acquisition

Myelin histology was obtained and processed from case P only as described in a previous study (Reveley et al., 2022). Briefly, the left hemisphere of case P was sectioned at 50 µm on a freezing microtome. We used the Gallyas silver stain (Gallyas, 1979) which stains individual axons at high resolution. Each section was digitally scanned at 0.88 μm resolution (×10 magnification) using a Zeiss Axioscan microscope slide scanner. Nine Gallyas stained sections were selected for nonlinear registration based on criteria of even staining and minimal damage in the previous study, and two of these were excluded from the present study due to a small, localized artefact in the dorsal part of the novel dMRI data described above.

### Histology processing

To estimate the anisotropy content of the myelin histology we employed the structure tensor (ST) method (Budde & Frank, 2012), as described in a previous study (Reveley et al., 2022). Briefly, the 0.88 μm per pixel resolution histology images were processed using the structure tensor as implemented in the “OrientationJ” ImageJ package using a local gaussian window σ = 85 pixels and image gradients computed with a cubic spline. Since the dMRI data contains diffusion information gathered over a 150μm cubic area, the histology structure tensor at each pixel was a distance weighted function of the structure tensors within a radius of 75 μm. For each pixel, the structure tensor coherence was computed from the ST eigenvalues.

### Histology Registration

The 75 µm MTR was manually rotated to the histology slice plane and the dMRI data registered to this using FLIRT as described above. Histology parameter maps of stain density and anisotropy at 75 µm resolution were then non-linearly registered to the MRI. We performed careful nonlinear alignment of seven local frontal and parietal ROI from the histology to the MRI. Stain intensity maps of the 75 µm maps were first binarized using a threshold adjustment in imageJ and then registered to their corresponding MRI slices to correct for shrinkage and deformation, first using manual 2D alignment in imageJ and then MIND MATLAB code83 (α = 0.1) as described in a previous study(Reveley et al., 2022).

### Data Analysis

#### Case M

We loaded the data into matlab and plotted it; describe figure by figure figure 3,4,5 Figure 3 A, B plotted in ImageJ. Figure 3 C, plotted in matlab – look up the segmentation in the code. Images are at 80um, correlations at 160 for figures 3,4, and 5.

#### Case P

Once rotated into the histology slice plane, each coronal slice of the dMRI parameter maps acquired for this study were converted to quantitative tif format and integrated into a software system written for a previous study on MRI histology comparisons (Reveley et al., 2022). Briefly, this system took advantage of the fact that cortex consists of vertically oriented tissue components to produce a matrix representation of the non-linearly registered dMRI and histology data whose columns were derived from data sampled along the vertical dMRI tractography and whose rows spanned the horizontal direction. Tractography lines were integrated through the 2d projected principal eigenvectors in each gray matter ROI. A vector of unique pixels whose coordinates intersected one streamline’s was acquired, each of which was then resampled to a fixed length using bilinear interpolation, to yield a simple matrix representation of the gray matter. This operation was invertible such that data in matrix form could be displayed in image space.

Our statistical analysis proceeded by parcellating these matrix representations of the nonlinearly registered histology and MRI data. The parcellation strategy served to separate out regions of potentially differing stain intensity, and to assess relationships between histology and MRI variables in different anatomical regions in a statistically independent manner. For each parcel we took the Pearson correlation between sample variables, in e.g., Figure 6 **C** and Figure 7 **A**. To assess the average laminar properties of histology or dMRI variables, we averaged the rows of the matrix representation and plotted the resulting vector, e.g., in Figure 6 **B** and Figure 7 **B.**

## Acknowledgments

The research was supported by the Intramural Research Program of the National Institute of Mental Health (ZIAMH002786 and ZICMH002899).

## Author Contributions

CR conceived the project, analyzed the data, wrote the paper. FQY acquired the data, advised on analyses, edited the paper. DAL advised on analyses, edited the paper.

## Data and Code Availability

Data and code will be made available on publication

## References

1. Andica, C., Kamagata, K., Hatano, T., Saito, Y., Ogaki, K., Hattori, N., & Aoki, S. (2020). MR Biomarkers of Degenerative Brain Disorders Derived From Diffusion Imaging. Journal of Magnetic Resonance Imaging, 52(6), 1620–1636. 10.1002/jmri.27019

2. Avants, B. B., Tustison, N. J., Song, G., Cook, P. A., Klein, A., & Gee, J. C. (2011). A reproducible evaluation of ANTs similarity metric performance in brain image registration. NeuroImage, 54(3), 2033–2044. 10.1016/j.neuroimage.2010.09.025

3. Avram, A. V., Saleem, K. S., Komlosh, M. E., Yen, C. C., Ye, F. Q., & Basser, P. J. (2022). High-resolution cortical MAP-MRI reveals areal borders and laminar substructures observed with histological staining: High-resolution cortical MAP-MRI. NeuroImage, 264(July), 119653. 10.1016/j.neuroimage.2022.119653

4. Balaram, P., & Kaas, J. H. (2014). Towards a unified scheme of cortical lamination for primary visual cortex across primates: Insights from NeuN and VGLUT2 immunoreactivity. Frontiers in Neuroanatomy, 8(AUG). 10.3389/fnana.2014.00081

5. Bester, M., Jensen, J. H., Babb, J. S., Tabesh, A., Miles, L., Herbert, J., Grossman, R. I., & Inglese, M. (2015). Non-Gaussian diffusion MRI of gray matter is associated with cognitive impairment in multiple sclerosis. Multiple Sclerosis, 21(7), 935–944. 10.1177/1352458514556295

6. Bock, N. A., Hashim, E., Kocharyan, A., & Silva, A. C. (2011). Visualizing myeloarchitecture with magnetic resonance imaging in primates. Annals of the New York Academy of Sciences, 1225(S1), E171–E181. 10.1111/j.1749-6632.2011.06000.x

7. Braitenberg, V. (1962). A note on myeloarchitectonics. Journal of Comparative Neurology, 118(2), 141–156. 10.1002/cne.901180202

8. Brodmann, K. (2006). Vergleichende lokalisationslehre der großhirnrinde in ihren prinzipien dargestellt auf grund des zellenbaues (1909)(English translation available in Garey, L.J. (2006). Brodmann’s: Localisation in the cerebral cortex. 3rd Edition. Springer US) (L. J. Garey (ed.); 3rd Editio). Springer US.

9. Budde, M. D., & Frank, J. A. (2012). Examining brain microstructure using structure tensor analysis of histological sections. NeuroImage, 63(1), 1–10. 10.1016/j.neuroimage.2012.06.042

10. Buxhoeveden, D. P., & Casanova, M. F. (2002). The minicolumn hypothesis in neuroscience. Brain, 125(5), 935–951. 10.1093/brain/awf110

11. Casagrande, V. A., & Kaas, J. H. (1994). The Afferent, Intrinsic, and Efferent Connections of Primary Visual Cortex in Primates. 10, 201–259. 10.1007/978-1-4757-9628-5_5

12. Daducci, A., Canales-Rodríguez, E. J., Zhang, H., Dyrby, T. B., Alexander, D. C., & Thiran, J. P. (2015). Accelerated Microstructure Imaging via Convex Optimization (AMICO) from diffusion MRI data. NeuroImage, 105, 32–44. 10.1016/j.neuroimage.2014.10.026

13. Edwards, L. J., Kirilina, E., Mohammadi, S., & Weiskopf, N. (2018). Microstructural imaging of human neocortex in vivo. NeuroImage, 182(October 2017), 184–206. 10.1016/j.neuroimage.2018.02.055

14. Falangola, M. F., Guilfoyle, D. N., Tabesh, A., Hui, E. S., Nie, X., Jensen, J. H., Gerum, S. V., Hu, C., Lafrancois, J., Collins, H. R., & Helpern, J. A. (2014). Histological correlation of diffusional kurtosis and white matter modeling metrics in cuprizone-induced corpus callosum demyelination. NMR in Biomedicine, 27(8), 948–957. 10.1002/nbm.3140

15. Fukutomi, H., Glasser, M. F., Zhang, H., Autio, J. A., Coalson, T. S., Okada, T., Togashi, K., Van Essen, D. C., & Hayashi, T. (2018). Neurite imaging reveals microstructural variations in human cerebral cortical gray matter. NeuroImage, 182(February), 488–499. 10.1016/j.neuroimage.2018.02.017

16. Gallyas, F. (1979). Silver staining of myelin by means of physical development. Neurological Research, 1(2), 203–209. http://www.ncbi.nlm.nih.gov/pubmed/95356

17. Glasser, M. F., Goyal, M. S., Preuss, T. M., Raichle, M. E., & Van Essen, D. C. (2014). Trends and properties of human cerebral cortex: Correlations with cortical myelin content. In NeuroImage (Vol. 93, pp. 165–175). Academic Press Inc. 10.1016/j.neuroimage.2013.03.060

18. Grossman, R. I., Gomori, J. M., Ramer, K. N., Lexa, F. J., & Schnall, M. D. (1994). Magnetization transfer: theory and clinical applications in neuroradiology. *Radiographics : A Review Publication of the Radiological Society of North America*, Inc, 14(2), 279–290. 10.1148/radiographics.14.2.8190954

19. Guglielmetti, C., Veraart, J., Roelant, E., Mai, Z., Daans, J., Van Audekerke, J., Naeyaert, M., Vanhoutte, G., Delgado y Palacios, R., Praet, J., Fieremans, E., Ponsaerts, P., Sijbers, J., Van der Linden, A., & Verhoye, M. (2016). Diffusion kurtosis imaging probes cortical alterations and white matter pathology following cuprizone induced demyelination and spontaneous remyelination. NeuroImage, 125, 363–377. 10.1016/j.neuroimage.2015.10.052

20. Heath, F., Hurley, S. A., Johansen-Berg, H., & Sampaio-Baptista, C. (2018). Advances in noninvasive myelin imaging. Developmental Neurobiology, 78(2), 136–151. 10.1002/dneu.22552

21. Henkelman, R. M., Stanisz, G. J., & Graham, S. J. (2001). Magnetization transfer in MRI: A review. NMR in Biomedicine, 14(2), 57–64. 10.1002/nbm.683

22. Henriques, R. N., Jespersen, S. N., Jones, D. K., & Veraart, J. (2021). Toward more robust and reproducible diffusion kurtosis imaging. Magnetic Resonance in Medicine, 86(3), 1600–1613. 10.1002/mrm.28730

23. Howard, A. F. D., Huszar, I. N., Cottaar, M., Daubney, G., Khrapitchev, A. A., Mars, R. B., Mollink, J., Roumazeilles, L., Scott, C., & Smart, A. (n.d.). The BigMac dataset: interconnecting MR signals with microstructure profiles in the cortex.

24. Jelescu, I. O., de Skowronski, A., Geffroy, F., Palombo, M., & Novikov, D. S. (2022). Neurite Exchange Imaging ((NEXI): A minimal model of diffusion in gray matter with inter-compartment water exchange. NeuroImage, 256. 10.1016/j.neuroimage.2022.119277

25. Jelescu, I. O., Zurek, M., Winters, K. V., Veraart, J., Rajaratnam, A., Kim, N. S., Babb, J. S., Shepherd, T. M., Novikov, D. S., Kim, S. G., & Fieremans, E. (2016). In vivo quantification of demyelination and recovery using compartment-specific diffusion MRI metrics validated by electron microscopy. NeuroImage, 132, 104–114. 10.1016/j.neuroimage.2016.02.004

26. Jensen, J. H., Helpern, J. A., Ramani, A., Lu, H., & Kaczynski, K. (2005). Diffusional kurtosis imaging: The quantification of non-Gaussian water diffusion by means of magnetic resonance imaging. Magnetic Resonance in Medicine, 53(6), 1432–1440. 10.1002/mrm.20508

27. Kamiya, K., Hori, M., & Aoki, S. (2020). NODDI in clinical research. Journal of Neuroscience Methods, 346(August), 108908. 10.1016/j.jneumeth.2020.108908

28. Kandel, E. R., Schwartz, J. H., Jessell, T. M., Siegelbaum, S. A., & Hudspeth, A. J. (2013). Principles of Neural Science (E. R. Kandel, J. H. Schwartz, T. M. Jessell, S. A. Siegelbaum, & A. J. Hudspeth (eds.); 5th ed.). McGraw Hill Professional. https://books.google.co.uk/books?id=s64z-LdAIsEC

29. Kelm, N. D., West, K. L., Carson, R. P., Gochberg, D. F., Ess, K. C., & Does, M. D. (2016). Evaluation of diffusion kurtosis imaging in ex vivo hypomyelinated mouse brains. NeuroImage, 124, 612–626. 10.1016/j.neuroimage.2015.09.028

30. Kirkcaldie, M. T. K., Dickson, T. C., King, C. E., Grasby, D., Riederer, B. M., & Vickers, J. C. (2002). Neurofilament triplet proteins are restricted to a subset of neurons in the rat neocortex. Journal of Chemical Neuroanatomy, 24(3), 163–171. 10.1016/S0891-0618(02)00043-1

31. Lazari, A., & Lipp, I. (2021). Can MRI measure myelin? Systematic review, qualitative assessment, and meta-analysis of studies validating microstructural imaging with myelin histology. NeuroImage, 230(September 2020), 117744. 10.1016/j.neuroimage.2021.117744

32. Lee, H. H., Papaioannou, A., Novikov, D. S., & Fieremans, E. (2020). In vivo observation and biophysical interpretation of time-dependent diffusion in human cortical gray matter. NeuroImage, 222(June), 117054. 10.1016/j.neuroimage.2020.117054

33. Liu, C., Ye, F. Q., Newman, J. D., Szczupak, D., Tian, X., Yen, C. C.-C., Majka, P., Daniel, G., Rosa, M. G. P., Leopold, D. A., & Silva, A. C. (2019). A resource for detailed 3D mapping of white matter pathways in the marmoset brain. *Nature Neuroscience*, (accepted. 10.1038/s41593-019-0575-0

34. Liu, C., Ye, F. Q., Newman, J. D., Szczupak, D., Tian, X., Yen, C. C. C., Majka, P., Glen, D., Rosa, M. G. P., Leopold, D. A., & Silva, A. C. (2020). A resource for the detailed 3D mapping of white matter pathways in the marmoset brain. Nature Neuroscience, 23(2), 271–280. 10.1038/s41593-019-0575-0

35. Liu, C., Ye, F. Q., Yen, C. C.-C. C., Newman, J. D., Glen, D., Leopold, D. A., Silva, A. C., Chern-Chyi Yen, C., Newman, J. D., Glen, D., Leopold, D. A., Silva, A. C., Yen, C. C.-C. C., Newman, J. D., Glen, D., Leopold, D. A., Silva, A. C., Chern-Chyi Yen, C., Newman, J. D., … Silva, A. C. (2018). A digital 3D atlas of the marmoset brain based on multi-modal MRI. NeuroImage, 169(July 2017), 106–116. 10.1016/j.neuroimage.2017.12.004

36. Mancini, M., Karakuzu, A., Cohen-Adad, J., Cercignani, M., Nichols, T. E., & Stikov, N. (2020). An interactive meta-analysis of MRI biomarkers of Myelin. ELife, 9, 1–23. 10.7554/eLife.61523

37. McKavanagh, R., Torso, M., Jenkinson, M., Kolasinski, J., Stagg, C. J., Esiri, M. M., McNab, J. A., Johansen-Berg, H., Miller, K. L., & Chance, S. A. (2019). Relating diffusion tensor imaging measurements to microstructural quantities in the cerebral cortex in multiple sclerosis. Human Brain Mapping, 40(15), 4417–4431. 10.1002/hbm.24711

38. Morell, P., & Quarles, R. H. (1999). The myelin sheath. *Basic Neurochemistry: Molecular*, Cellular and Medical Aspects, 6.

39. Novikov, D. S. (2021). The present and the future of microstructure MRI: From a paradigm shift to normal science. Journal of Neuroscience Methods, 351, 108947. 10.1016/j.jneumeth.2020.108947

40. Ouyang, M., Jeon, T., Sotiras, A., Peng, Q., Mishra, V., Halovanic, C., Chen, M., Chalak, L., Rollins, N., Roberts, T. P. L., Davatzikos, C., & Huang, H. (2019). Differential cortical microstructural maturation in the preterm human brain with diffusion kurtosis and tensor imaging. Proceedings of the National Academy of Sciences of the United States of America, 116(10), 4681–4688. 10.1073/pnas.1812156116

41. Özarslan, E., Koay, C. G., Shepherd, T. M., Komlosh, M. E., İrfanoğlu, M. O., Pierpaoli, C., Basser, P. J., Irfanoǧlu, M. O., Pierpaoli, C., Basser, P. J., İrfanoğlu, M. O., Pierpaoli, C., & Basser, P. J. (2013). Mean apparent propagator (MAP) MRI: A novel diffusion imaging method for mapping tissue microstructure. NeuroImage, 78, 16–32. 10.1016/j.neuroimage.2013.04.016

42. Paxinos, G., Watson, C., Petrides, M., Rosa, M., & Tokuno, H. (2013). The Marmoset Brain in Stereotaxic Coordinates. https://espace.curtin.edu.au/handle/20.500.11937/40725

43. Paydar, A., Fieremans, E., Nwankwo, J. I., Lazar, M., Sheth, H. D., Adisetiyo, V., Helpern, J. A., Jensen, J. H., & Milla, S. S. (2014). Diffusional kurtosis imaging of the developing brain. American Journal of Neuroradiology, 35(4), 808–814. 10.3174/ajnr.A3764

44. Peters, A. (2010). The Morphology of Minicolumns. The Neurochemical Basis of Autism: From Molecules to Minicolumns, 1–295. 10.1007/978-1-4419-1272-5

45. Peters, A., Manuel Cifuentes, J., & Sethares, C. (1997). The organization of pyramidal cells in area 18 of the rhesus monkey. Cerebral Cortex, 7(5), 405–421. 10.1093/cercor/7.5.405

46. Peters, A., & Sethares, C. (1996). Myelinated axons and the pyramidal cell modules in monkey primary visual cortex. Journal of Comparative Neurology, 365(2), 232–255. 10.1002/(SICI)1096-9861(19960205)365:2<232::AID-CNE3>3.0.CO;2-6

47. Pierpaoli, C., Walker, L., Irfanoglu, M., Barnett, A., Basser, P., Chang, L., Koay, C., Pajevic, S., Rohde, G., Sarlls, J., & others. (2010). TORTOISE: an integrated software package for processing of diffusion MRI data.

48. Praet, J., Manyakov, N. V., Muchene, L., Mai, Z., Terzopoulos, V., De Backer, S., Torremans, A., Guns, P. J., Van De Casteele, T., Bottelbergs, A., Van Broeck, B., Sijbers, J., Smeets, D., Shkedy, Z., Bijnens, L., Pemberton, D. J., Schmidt, M. E., Van Der Linden, A., & Verhoye, M. (2018). Diffusion kurtosis imaging allows the early detection and longitudinal follow-up of amyloid-β-induced pathology. Alzheimer’s Research and Therapy, 10(1). 10.1186/s13195-017-0329-8

49. Reina, M. A., Arriazu Navarro, R., & Durán Mateos, E. M. (2015). Ultrastructure of Myelinated and Unmyelinated Axons. In Atlas of Functional Anatomy for Regional Anesthesia and Pain Medicine (pp. 3–18). Springer International Publishing. 10.1007/978-3-319-09522-6_1

50. Reveley, C., Ye, F. Q., Mars, R. B., Matrov, D., Chudasama, Y., & Leopold, D. A. (2022). Diffusion MRI anisotropy in the cerebral cortex is determined by unmyelinated tissue features. Nature Communications, 13(1). 10.1038/s41467-022-34328-z

51. Schneider, T., & Wheeler-Kingshott, C. A. M. (2014). Q-Space Imaging: A Model-Free Approach. In Quantitative MRI of the Spinal Cord (pp. 146–155). Elsevier Inc. 10.1016/B978-0-12-396973-6.00010-1

52. Torso, M., Bozzali, M., Zamboni, G., Jenkinson, M., & Chance, S. A. (2021). Detection of Alzheimer’s Disease using cortical diffusion tensor imaging. Human Brain Mapping, 42(4), 967–977. 10.1002/hbm.25271

53. Torso, M., Ridgway, G. R., Valotti, M., Hardingham, I., Chance, S. A., Brewer, J., Lopez, O., Hyman, B.,Grabowski, T., Sano, M., Chui, H., Albert, M., Morris, J., Kaye, J., Wisniewski, T., Small, S., Trojanowski, J., DeCarli, C., Saykin, A., … Leverenz, J. (2023). In vivo cortical diffusion imaging relates to Alzheimer’s disease neuropathology. Alzheimer’s Research and Therapy, 15(1), 1–15. 10.1186/s13195-023-01309-3

54. Vavasour, I. M., Laule, C., Li, D. K. B., Traboulsee, A. L., & MacKay, A. L. (2011). Is the magnetization transfer ratio a marker for myelin in multiple sclerosis? Journal of Magnetic Resonance Imaging, 33(3), 710–718. 10.1002/jmri.22441

55. Vogt, C., & Vogt, O. (1919). Allgemeinere Ergebnisse unserer Hirnforschung. J Psychol Neurol (Leipz), 25, 279–461. http://ci.nii.ac.jp/naid/10024136586/en/

56. Wang, F., Dong, Z., Tian, Q., Liao, C., Fan, Q., Hoge, W. S., Keil, B., Polimeni, J. R., Wald, L. L., Huang, S. Y., & Setsompop, K. (2021). In vivo human whole-brain Connectom diffusion MRI dataset at 760 µm isotropic resolution. Scientific Data, 8(1), 1–12. 10.1038/s41597-021-00904-z

57. Weston, P. S. J., Poole, T., Nicholas, J. M., Toussaint, N., Simpson, I. J. A., Modat, M., Ryan, N. S., Liang, Y., Rossor, M. N., Schott, J. M., Ourselin, S., Zhang, H., & Fox, N. C. (2020). Measuring cortical mean diffusivity to assess early microstructural cortical change in presymptomatic familial Alzheimer’s disease. Alzheimer’s Research and Therapy, 12(1), 1–10. 10.1186/s13195-020-00679-2

58. Wolff, S. D., & Balaban, R. S. (1989). Magnetization transfer contrast (MTC) and tissue water proton relaxation in vivo. Magnetic Resonance in Medicine, 10(1), 135–144. 10.1002/mrm.1910100113

59. Woolrich, M. W., Jbabdi, S., Patenaude, B., Chappell, M., Makni, S., Behrens, T., Beckmann, C., Jenkinson, M., & Smith, S. M. (2009). Bayesian analysis of neuroimaging data in FSL. NeuroImage, *45*(1, Supplement 1), S173–S186. doi:10.1016/j.neuroimage.2008.10.055

60. Wu, E. X., & Cheung, M. M. (2010). MR diffusion kurtosis imaging for neural tissue characterization. NMR in Biomedicine, 23(7), 836–848. 10.1002/nbm.1506

61. Yoshida, M., Hori, M., Yokoyama, K., Fukunaga, I., Suzuki, M., Kamagata, K., Shimoji, K., Nakanishi, A.,Hattori, N., Masutani, Y., & Aoki, S. (2013). Diffusional kurtosis imaging of normal-appearing white matter in multiple sclerosis: Preliminary clinical experience. Japanese Journal of Radiology, 31(1), 50–55. 10.1007/s11604-012-0147-7

62. Zhang, H., Schneider, T., Wheeler-Kingshott, C. A., & Alexander, D. C. (2012). NODDI: Practical in vivo neurite orientation dispersion and density imaging of the human brain. NeuroImage, 61(4), 1000–1016. 10.1016/j.neuroimage.2012.03.072

63. Zhu, T., Peng, Q., Ouyang, A., & Huang, H. (2021). Neuroanatomical underpinning of diffusion kurtosis measurements in the cerebral cortex of healthy macaque brains. Magnetic Resonance in Medicine, 85(4), 1895–1908. 10.1002/mrm.28548

64. Zhuo, J., & Gullapalli, R. P. (2020). Diffusion Kurtosis Imaging. In M. Mannil & S. F.-X. Winklhofer (Eds.), Neuroimaging Techniques in Clinical Practice: Physical Concepts and Clinical Applications (pp. 215–228). Springer International Publishing. 10.1007/978-3-030-48419-4_15

