## Supplementary material for "Diffusion kurtosis MRI tracks gray matter myelin content in the primate cerebral cortex": SI Text and Figures

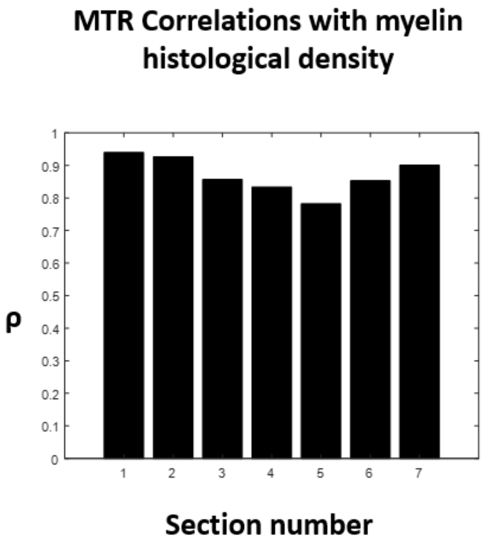


*SI Figure 1 Pearson correlation coefficients for pixelwise correlations between myelin stain density and MTR across 7 non-linearly registered sections.*

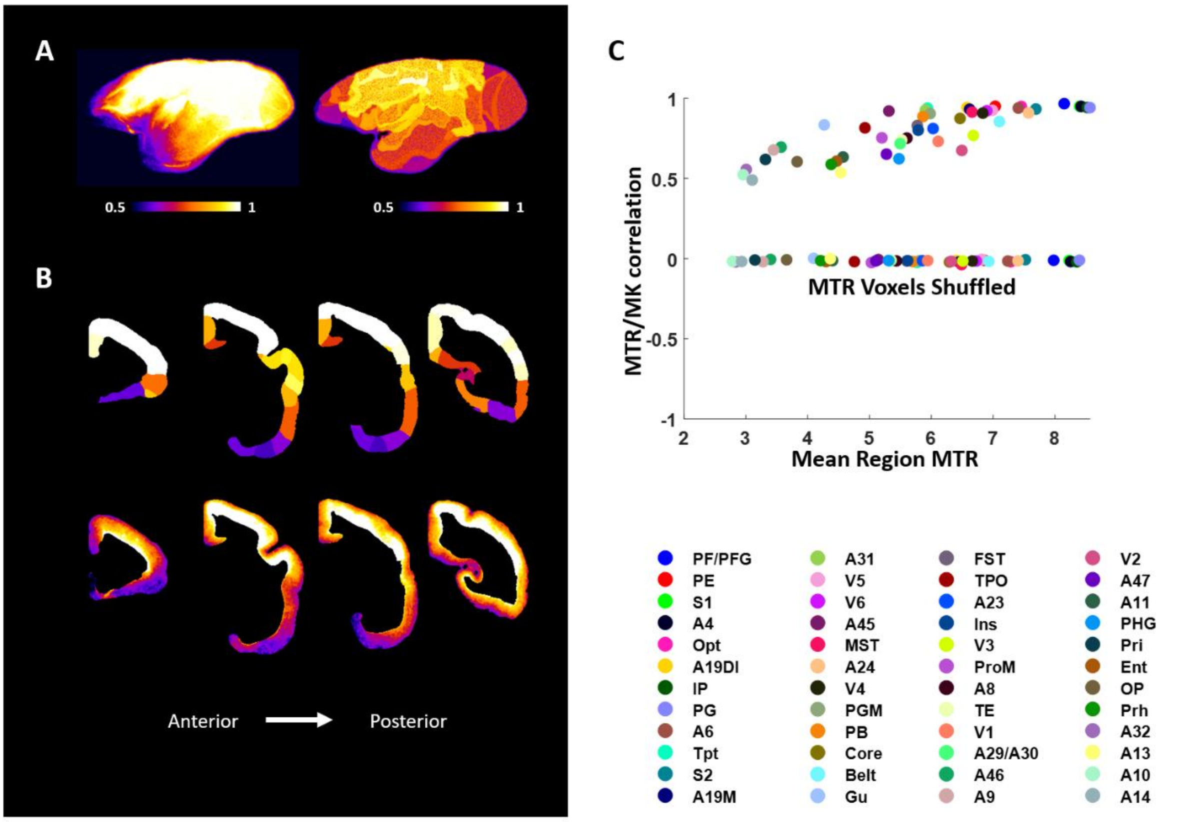


*SI Figure 2* ***Strength of correlation between MTR and MK is linked to tissue myelin levels A:*** *Surface maps showing MTR intensity (left) and the strength of correlation between MTR and MK (right); the parameters show a similar spatial distribution in the marmoset cortex.* ***B:*** *Select coronal sections of MK/MTR correlations (top row) and MTR (bottom row) at 80 µm resolution.* ***C:*** *Correlations of MK and MTR at 80 µm resolution for each cortical region. Pearson correlation coefficients between MK and MTR for each marmoset cortical region at 80 µm resolution as plotted against the mean MTR value for each region. Key shows regions delineated in the medium granularity cortical atlas of the marmoset brain mapping project* (Liu et al., 2018, 2019),

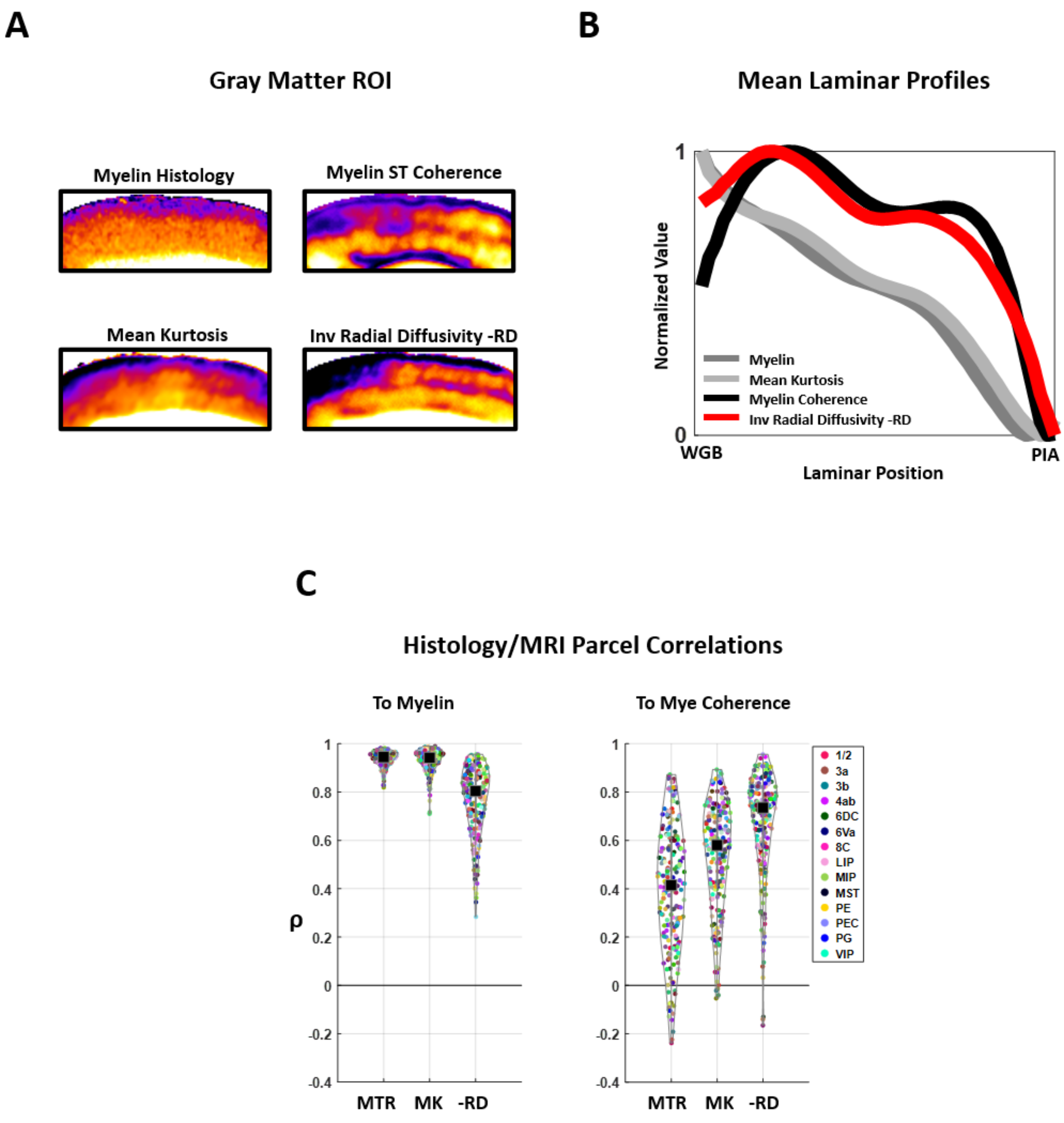


*SI Figure 3* ***Differences in the relationships of RD and MK to the density of histological myelin, and its organization assessed by structure tensor (ST) coherence (see Methods) A:*** *Representative ROI of histological myelin, histological myelin ST Coherence, MK and -RD.* ***B:*** *mean laminar profiles of Histological Myelin, MK, MTR and histological myelin coherence.* ***C:*** *parcel-wise correlations of MRI parameters to histological myelin and myelin coherence (see* ***Methods****).*

### *SI Text 1*

We found that mean kurtosis and radial diffusivity differed in their relationship to cortical myelin. Although MK very closely tracked myelin levels in the gray matter (**Figure 6**, **Figure 7**), it was unaffected by the orientations of the axons, whereas radial diffusivity was affected by both of these tissue features (**SI Figure 4**). To assess the orientation content of the myelinated fibers, we obtained a structure tensor derived measure of axonal orientation variation (ST Coherence) from the histology (see **Methods**). ST coherence is insensitive to the density of axons but provides a scalar index of how dispersed they are, with high values indicating low dispersion.

We found that -RD bore a strong resemblance to the spatial pattern of myelin ST coherence, while MK matches the pattern of myelin density (**SI Figure 4 A**). Taking the average laminar profile of the dMRI and histology data, we found that while both myelin levels and mean kurtosis showed a monotonic decline from deep to superficial layers, -RD showed a more complex curve that was congruent with the degree of alignment of the stained axons as measured by ST Coherence (**SI Figure 4 B**). When we analyzed the correlations between MK and -RD for each parcel of cortex across 7 histological sections (see **Methods**) we found that while -RD did have a significant mean correlation to myelin levels (**SI Figure 4 C, left panel**), it was lower than that of MK. In contrast, -RD showed a higher correlation to myelin ST coherence than either MTR or MK.

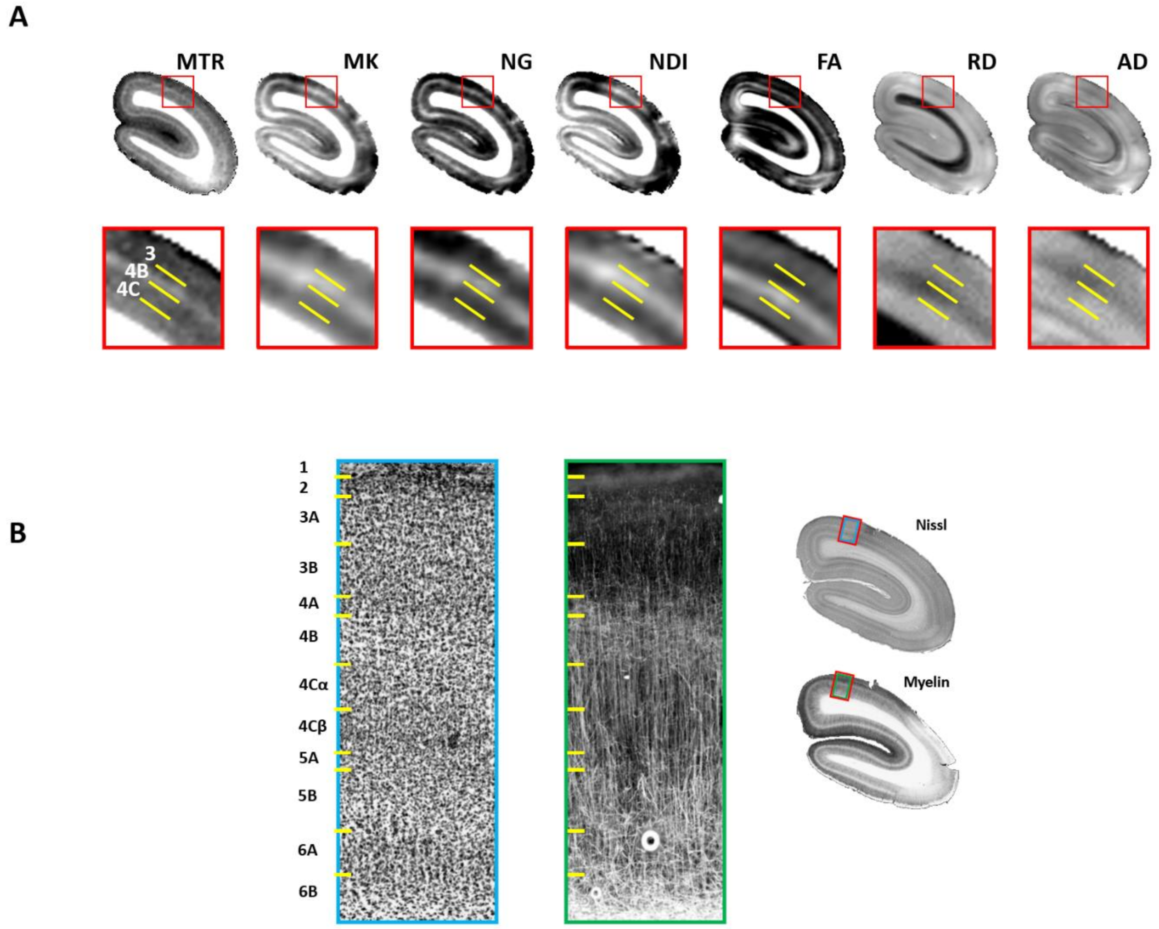


*SI Figure 4* ***MTR, MK, NG, NDI reveal the same laminar properties of myelin in primary visual cortex, while FA, RD and AD reflect more complex tissue properties including non-myelinated components A:*** *MTR, MK, NG, NDI, FA RD and AD parameter maps expose different cortical layers and sublayers. In particular, the high myelin Gennari band of layer 4B is exposed as a high intensity band of MTR, MK, NG and NDI, but a low intensity band of FA. In contrast, Layer 4C is shown as a low intensity band of MTR, MK and NDI but a high intensity band of FA, related to decreased RD and elevated AD .* ***B:*** *Nissl and myelin histology from the same subject matched to the MRI shown in A. See text for discussion.*

*SI Text 2*

We investigated the laminar dMRI properties of primary visual cortex (**V1**) with a detailed comparison of MRI to myelin and Nissl histology. Anatomically, V1 exhibits strongly contrasting layers with sharp laminar boundaries and wide variation in myelin levels, myeloarchitecture and cytoarchitecture between layers. V1 anatomy was reflected both in great detail and with high definition by our dMRI data; differences in laminar myelin levels were reflected in MK, NG and NDI intensity as well as in the MTR, while many other laminar features were reflected in distinctive patterns of anisotropy and diffusivity (**SI Figure 5**). In particular, we were able to clearly distinguish sublayers of layer 4.

In primates, layer 4B comprises a prominent band of myelinated fibers known as the stria of Gennari. A band of very high intensity in MTR and MK exposed layer 4B in our marmoset data (**SI Figure 5 A**), however, this layer had almost negligible FA (**SI Figure 5 A**). The abundant myelinated axons in 4B were revealed by histology to have a matted structure (**SI Figure 5 B**). This lack of spatial coherence likely explains the very low FA in this layer. Although the myelin content of the Gennari band is high, radial diffusivity remains high. Axial diffusivity is relatively low in layer 4B, which may relate to the relative paucity of apical dendrites in this layer of V1 (Peters, 2010). Together these diffusivity values produce low anisotropy.

Moving deeper, we found in the histology that layer 4C showed much lower myelin levels than 4B and a high density of small granule cells (**SI Figure 5 B**). As expected from the low myelin content (**SI Figure 5 B**) MTR, MK NG and NDI intensities were all diminished in layer 4C; however, we found FA to be greatly elevated. Radial diffusivity was depressed in layer 4C, likely owing to the fact that myelinated axons in this layer are organized in parallel to each other. Axial diffusivity was elevated, likely due to a greater number of coherently organized dendrites and unmyelinated axon terminals (Peters, 2010).
